# Antimicrobial wound care in an ant society

**DOI:** 10.1101/2022.04.26.489514

**Authors:** Erik. T. Frank, Lucie Kesner, Joanito Liberti, Quentin Helleu, Adria C. LeBoeuf, Andrei Dascalu, Douglas B. Sponsler, Fumika Azuma, Evan P. Economo, Patrice Waridel, Philipp Engel, Thomas Schmitt, Laurent Keller

## Abstract

Infected wounds pose a major mortality risk in animals^1,2^. Injuries are common in the ant *Megaponera analis*, which raids pugnacious prey^3,4^. Here we show that *M. analis* can determine when wounds are infected and treat them accordingly. By applying a variety of antimicrobial compounds and proteins secreted from the metapleural gland to infected wounds, workers reduce the mortality of infected individuals by 90%. Chemical analyses showed that wound infection is associated with specific changes in the cuticular hydrocarbon profile, thereby likely allowing nestmates to diagnose the infection state of injured individuals and apply the appropriate antimicrobial treatment. This study demonstrates that the targeted use of antimicrobials to treat infected wounds, previously thought to be a uniquely human behavior, has evolved in insect societies as well.

---

Infected wounds are a major mortality risk for animals^1,2^, but the identification and medicinal treatment of infected wounds has thus far been considered a uniquely human behavior. While several mammals have been shown to lick wounds and apply saliva^1,2^, the efficacy of this behavior remains largely unknown and occurs irrespective of the state of the wound. Workers of the predatory ant *Megaponera analis* have been shown to care for the injuries of nestmates^3,4^, which are common because this ant feeds exclusively on pugnacious termite species. As many as 22% of the foragers engaging in raids attacking termites have one or two missing legs^3^. Injured workers are carried back to the nest where other workers treat their wounds^4^. When the wounds of injured workers are not treated by nestmates, 90% of the injured workers die within 24 hours after injury^4^, but the mechanisms behind these treatments are unknown.

To investigate whether the high mortality of injured individuals is due to infection by pathogens, we collected soil from the natural environment and applied it to the wounds of experimentally injured workers (i.e., a sterile cut in the middle of the femur on the hind leg of an otherwise healthy ant). After 2 hours, injured ants exposed to the soil (hereafter referred to as “infected ants”) had ten times higher bacterial loads in the thorax than similarly injured individuals exposed to sterile phosphate buffered saline (PBS, hereafter referred to as “sterile ants”; least square means: *t*=-3.08; *P*=0.01; Fig. 1a, Supplemental Table 1). After 11 hours, bacterial load further increased in infected ants (least square means: *t*=-3.66; *P*=0.003), while there was no such increase in sterile ants (least square means: *t*=-1.54; *P*=0.26; Fig. 1a). As a result, after 11 hours the bacterial load was 100 times higher in infected than sterile ants (least square means: *t*=-5.21; *P*<0.001; Fig. 1a). A microbiome analysis further revealed major differences in bacterial species composition and abundance between the two groups of ants (ADONIS: *F*=17.45; *R*^2^=0.31; *P*<0.001; Fig. 1b & Extended Data Fig. 1a, b), with three potentially pathogenic bacterial genera increasing in absolute abundance in the thorax of infected ants at both time-points: *Klebsiella, Pseudomonas* and *Burkholderia* (Fig. 1b, Supplemental Table 2). These differences in bacterial composition and abundance between infected and sterile ants were associated with important differences in survival probability, with survival being seven times lower for infected ants (least square means: *Z*=-4.246; *P*<0.001; Fig. 1c, Extended Data Fig. 2a & Supplemental Table 3).

**Fig. 1.**
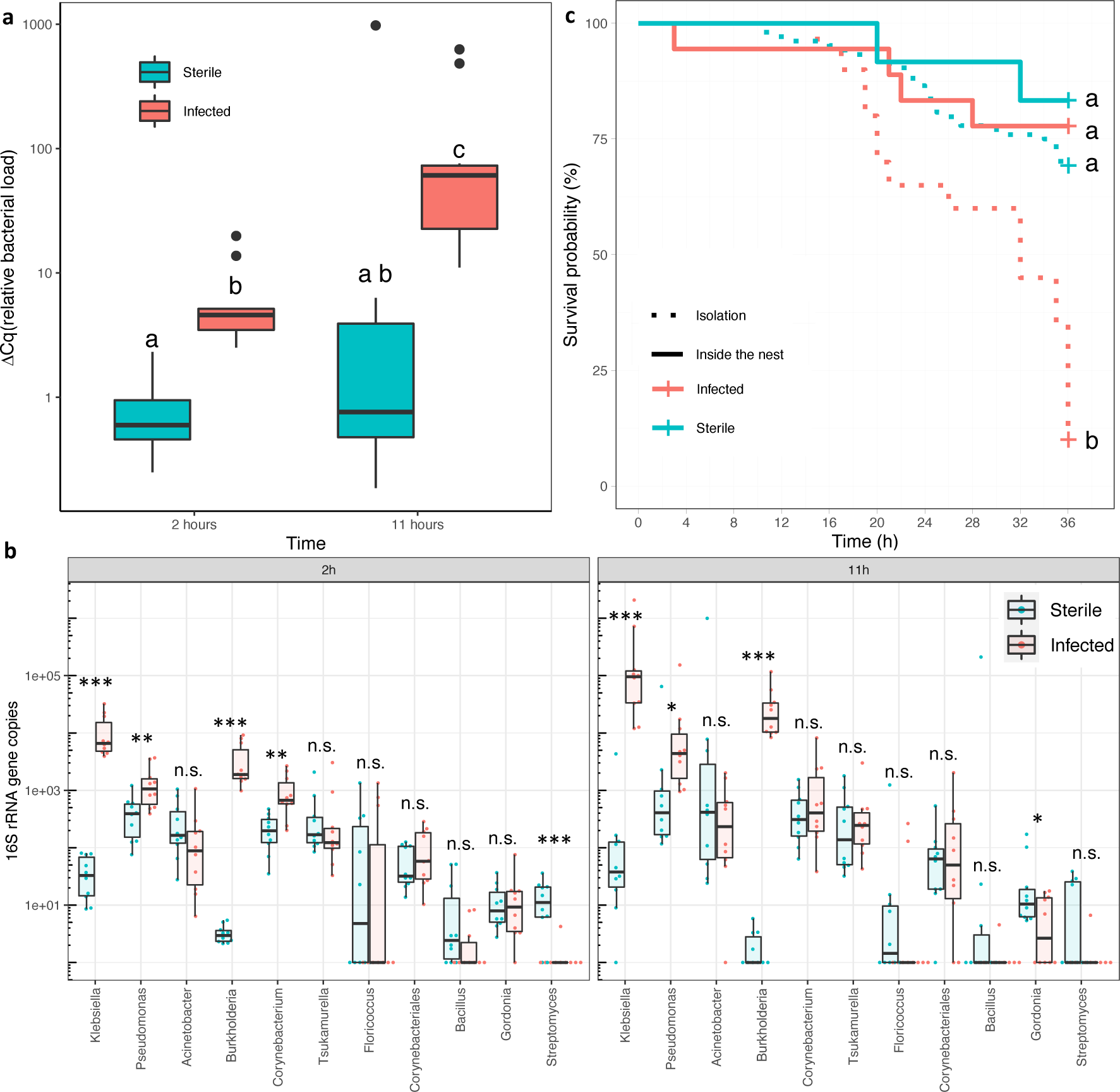
Lethal effects and diversity of soil pathogens. **(a)** Relative 16S rRNA gene copies (bacterial load 1′Cq) for individuals whose wounds were exposed to a sterile PBS solution (Sterile), or soil pathogens diluted in PBS (Infected, OD=0.1), 2 and 11 hours after exposure (see Supplemental Table 1 for statistical results). *n*=10 per boxplot. Significant differences (*P*<0.05) are shown with different letters. **(b)** Absolute 16S rRNA gene copy numbers summarized at the genus-level for the 18 amplicon sequence variants (ASV) that had at least 1% relative abundance in five of the 40 analyzed ants across the experiment (*n*=10 per boxplot). Significance is shown for pairwise comparisons between sterile and infected ants: ***=*P*<0.001; **=*P*<0.01; *=*P*<0.05; n.s.= not significant (*P*>0.05). Detailed statistical results in Supplemental Table 2 **(c)** Kaplan – Meier cumulative survival rates of workers in isolation (dotted line) or inside the nest (solid line) whose wounds were exposed the same way as in Fig. 1a (infected or sterile). Significant differences (*P*<0.05) are indicated with different letters (detailed statistical results in Extended Data Fig. 2a and Supplemental Table 3).

**Fig. 2.**
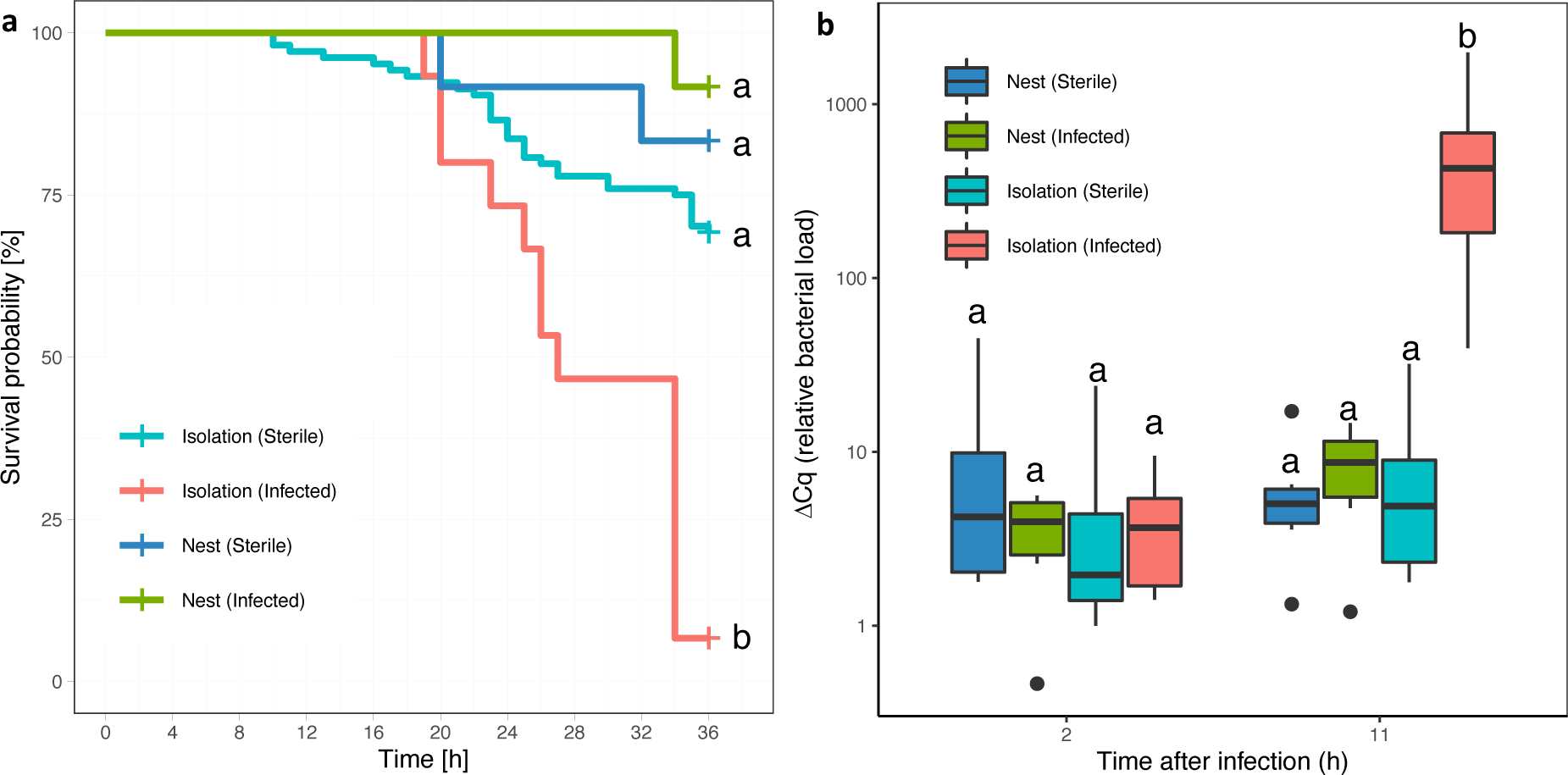
Survival probability and pathogen load of sterile and infected ants. **(a)** Kaplan – Meier cumulative survival rates of workers in isolation (dotted line) or inside the nest (solid line) whose wounds were exposed to *P. aeruginosa* diluted in PBS (Infected, OD=0.05) or a sterile PBS solution (Sterile). Detailed statistical results in Extended Data Fig. 2c and Supplemental Table 3. **(b)** relative bacterial load (1′Cq) of *Pseudomonas* at two different time points (2h and 11h) for ants in isolation or inside the nest with wounds treated the same way as in Fig. 2a (Infected or Sterile). *n*=6 per boxplot, significant differences (*P*<0.05) are shown with different letters (Supplemental Table 4).

To test if wound care by nestmates could reduce the mortality of infected ants, we either placed infected and sterile ants in their colony or kept them in isolation. The mortality after 36 hours was much lower for infected ants kept with their nestmates (22%) than for infected ants kept in isolation (90%, least square means: *Z*=-2.759; *P*=0.02; Fig. 1c). By contrast, there was no significant difference between the mortality of sterile ants kept with their nestmates or in isolation (least square means: *Z*=1.04; *P*=0.89; Fig. 1c). The mortality of infected ants was also not significantly different than the mortality of sterile ants when these individuals were kept with their nestmates (least square means: *Z*=-0.630; *P*=1; Fig. 1c). Overall, these data demonstrate that *M. analis* workers are capable of effectively treating wounds that have been exposed to soil pathogens.

By culturing the soil medium on agar plates, we were able to isolate three potential pathogens (the endosymbiotic bacterium *Burkholderia* sp. and its fungal host *Rhizopus microsporus*, and the bacterium *Pseudomonas aeruginosa*, Fig. 1b, Extended Data Fig. 1c, d). While the application of *B.* sp. and *R. microsporus* (separately or together) on wounds did not significantly decrease survival (Extended Data Fig. 2b & 3), the application of *P. aeruginosa* (OD=0.05), a bacterium widespread in various environments^5^, caused a mortality of 93% within 36 hours (Fig. 2a). Since the treatment with only *P. aeruginosa* showed a similar mortality as the treatment with all soil pathogens (90%, Extended Data Fig. 3), we only used *P. aeruginosa* in subsequent infection assays to better control pathogen load.

**Fig. 3.**
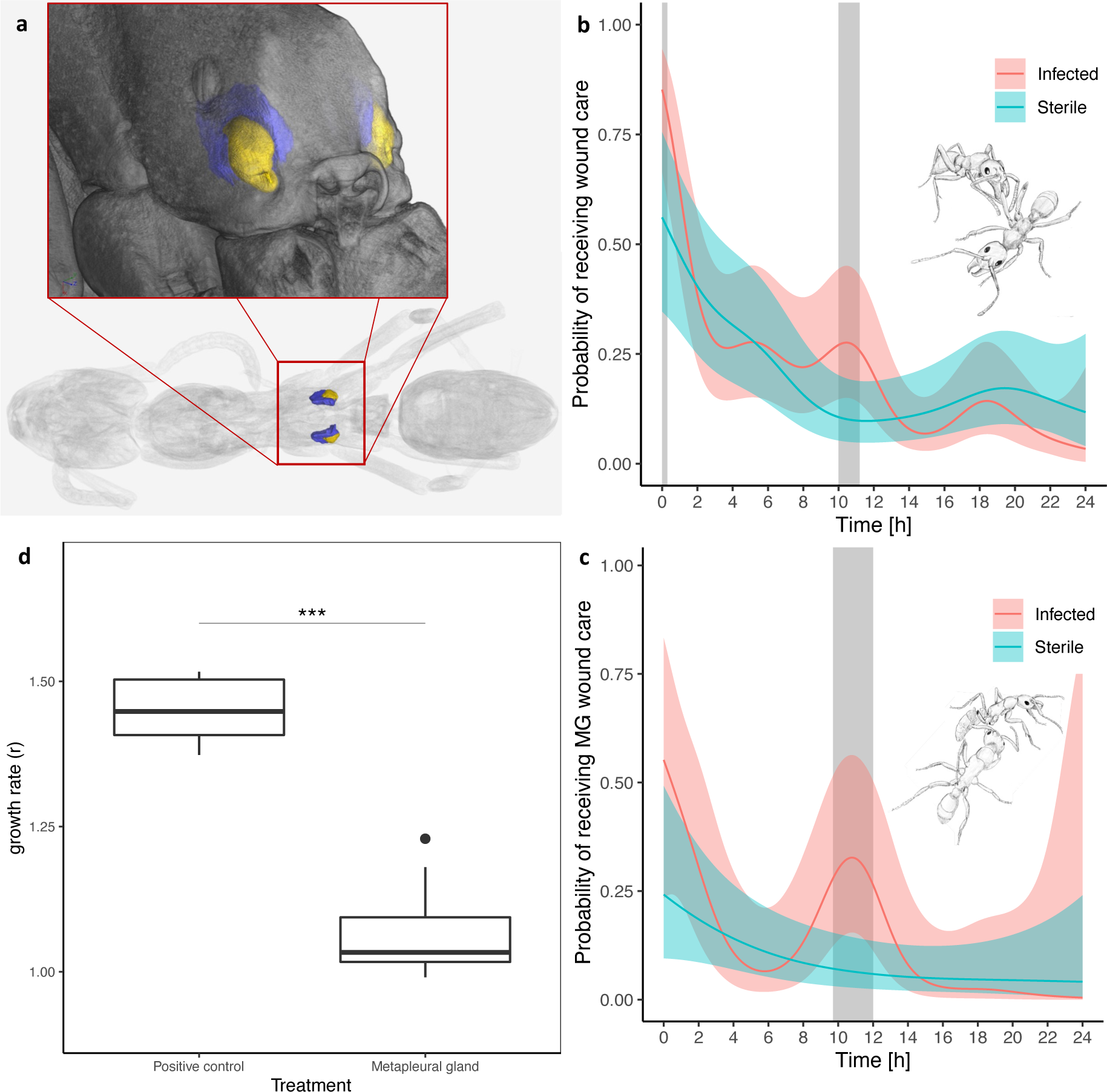
Use and efficacy of the metapleural gland (MG) secretions during wound care. **(a)** Micro CT scan showing the location of the MG. Blue: secretory cells; yellow: atrium. **(b)** Probability of receiving wound care over 24 hours fitted with a hierarchical generalized (binomial) additive model (HGAM); shaded bars indicate periods during which the probability of receiving care was significantly (P<0.05) higher for infected (n=6) than sterile (n=6) individuals. **(c)** Probability of receiving antimicrobial wound care with MG secretions (the same ants as in Fig. 3b), modeled with identical HGAM specifications. **(d)** Bacterial growth assay for *P. aeruginosa* either in LB broth (positive control, *n*=6) or LB broth with MG secretions (Metapleural gland *n*=9). Mann-Whitney U test: *W*=54, *P*<0.001.

Like the experiments where the soil was applied to the wound, the presence of nestmates was also effective in decreasing the mortality of injured workers exposed to a known concentration of *P. aeruginosa* (OD=0.05). While the mortality of infected ants kept in isolation was 93%, mortality of infected ants that had been returned to their nestmates was only 8% (least square means: *Z*=-2.94; *P*=0.01; Fig. 2a, Extended Data Fig. 2c & Supplemental Table 4). There were major differences in the increase in *Pseudomonas* load after injury between ants kept with or without their nestmates (Fig. 2b & Supplemental Table 5). The bacterial load of infected ants kept with their nestmates did not increase significantly from 2 and 11 hours after injury (least square means: *t*=-0.037, *P*=1; Fig. 2b). By contrast, there was a 100-fold increase in bacterial load for infected ants kept in isolation (least square means: *t*=-4.832, *P*<0.001; Fig. 2b).

To study the proximate mechanisms reducing mortality of infected ants when they are returned to their nestmates, we introduced injured ants (with sterile and infected wounds) to their nestmates and filmed them for 24 hours. We observed that workers treated the injury of infected ants by depositing secretions produced by the metapleural gland (MG), which is located at the back of the thorax (Fig. 3a). The MG secretions, which have antimicrobial properties^6-10^, were applied in 10.5% of the wound care interactions (43 out of 411). Before applying MG secretions, the nursing ant always groomed the wound first (i.e., “licking” the wound with their mouthparts). Nursing ants then collected the secretions either from their own MGs (Supplemental Movie 1), by reaching back to the opening of the gland with their front legs to collect the secretion and then licking their front legs to accumulate it in their mouth, as described in other species^8^, or from the MG of the injured ant itself, by licking directly into the gland’s opening (Supplemental Movie 2). Wound care with MG secretions lasted significantly longer (85±53s) than wound care without MG secretions (53±36s; t-test: *t*=-3.09, *P*=0.003; Extended Data Fig. 4). Remarkably, workers were able to discriminate between infected and sterile ants. Wound care treatment was provided more often to infected ants upon initial introduction to the nest and again 10- and 11-hours post-introduction (hierarchical generalized additive model: P<0.05; Fig. 3b). Moreover, MG secretions were deposited significantly more often on wounds of infected than sterile ants between 10 and 12 hours after infection (hierarchical generalized additive model: P<0.05; Fig. 3c). When treatment with MG secretions was prevented (through plugging the MG opening of all ants in the sub-colony), mortality of infected ants reached 100% within 36h compared to only 33% when workers had access to the MG secretions (least square means: *Z*=2.99; *P*=0.039; Fig. 4; Extended Data Fig. 5 & Supplemental Table 6). By contrast, the plugging of the MG opening had no significant effect on the survival of sterile ants (least square means: Z=0.02; *P*=1; Fig. 4, fig. S5 & table S6).

**Fig. 4.**
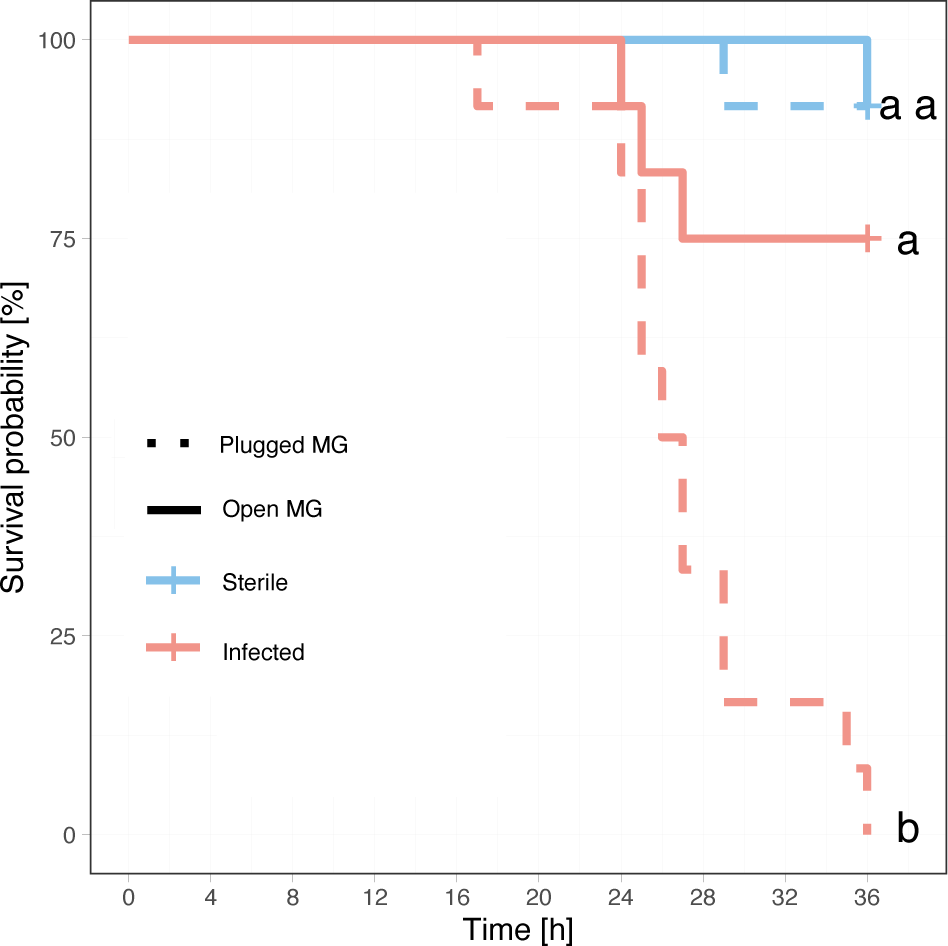
Effect of MG secretions on survival of sterile and infected ants. Kaplan – Meier cumulative survival rates of workers inside sub-colonies with a plugged MG opening (dotted line) or with an unmanipulated MG opening (solid line) whose wounds were exposed to a sterile PBS solution (Sterile *n*=12) or *P. aeruginosa* diluted (OD=0.05) in PBS (Infected *n*=12). Detailed statistical results in Extended Data Fig. 2d and Supplemental Table 5. Data for ants in isolation in Extended Data Fig 5.

Because cuticular hydrocarbons (CHCs) are known to be frequently used as a source of information in ants^11^, we investigated whether infected ants could signal their injured state through changes in the profile. Immediately after injury, infected ants did not differ from sterile ants in their CHC profile (ADONIS: *R*^2^=0.13; *F*=0.13; *P*=0.96; Supplemental Table 7 & 8). This profile changed in both types of ants during the two hours after injury (Sterile ants: ADONIS: *R*^2^=0.15; *F*=2.74; *P*=0.03; Infected ants: ADONIS: *R*^2^=0.13; *F*=2.42; *P*=0.05), converging towards a similar profile for both types of ants (ADONIS: *R*^2^=0.008; *F*=0.17; *P*=0.87). Thereafter, the CHC profile of infected ants remained unchanged until 11 hours after injury (ADONIS: *R*^2^=0.063; *F*=1.48; *P*=0.13), while the CHC profile of sterile ants changed significantly (ADONIS: *R*^2^=0.14; *F*=3.54; *P*=0.04), thereby becoming significantly different from the CHC profile of infected ants (ADONIS: *R*^2^=0.19; *F*=5.4; *P*=0.007). Eleven hours after injury, the CHC profile of sterile ants had converged again to the profile at 0h (ADONIS: *R*^2^=0.03; *F*=0.53; *P*=0.45), while the CHC profile of injured ants remained significantly different (ADONIS: *R*^2^=0.19; *F*=3.82; *P*=0.008).

Consistent with the idea that changes in CHC profiles could provide information on the health status of ants^12^, the observed differences in the CHC profiles mostly stemmed from differences in the relative abundance of alkadienes (Extended Data Fig. 6, Supplemental Table 9), which are among the CHC compounds most relevant for communication in social insects^13^. These changes in the CHC profile are generally regulated by differentially expressed genes in the fat body^14^. To identify the genes likely responsible for the observed CHC changes between infected and sterile ants, we conducted transcriptomic analyses of the fat bodies of the same individuals. A total of 18 genes related to CHC synthesis were differentially expressed between infected and sterile ants 11 hours after exposure to *P. aeruginosa* (17 genes out of 378 that were differentially expressed were immune genes; Extended Data Fig. 7a, Supplemental Table 10 & 11), while only two genes related to CHC synthesis were differentially expressed two hours after infection (in addition to 7 immune genes out of 164; Extended Data Fig. 6b, Supplemental Table 10 & 11).

To quantify the capabilities of MG secretions to inhibit bacterial growth, we conducted antimicrobial assays. The growth of *P. aeruginosa* was reduced by >25% when MG secretions were included in a lysogeny broth (LB) solution compared to a control LB solution without the MG secretions (Mann-Whitney U test: *W*=54, *P*<0.001, Fig. 3d).

Since *P. aeruginosa* has repeatedly developed antimicrobial resistance^15^ and because most antimicrobial compounds found in animal saliva are unable to inhibit the growth of *P. aeruginosa*^16^, we conducted proteomic and chemical analyses of the MG secretions. The proteomic analysis revealed 41 proteins (Extended Data Fig. 8 & Supplemental Table 12), 15 of which showed molecular similarity to toxins, which often have antimicrobial properties^17^. Five proteins had orthologs known to have antimicrobial activity (e.g., lysozyme, hemocytes, 2 MRJP1-like proteins) and three with melanization, a process implicated in wound healing in insects^18,19^. There was no clear function for nine proteins including the most abundant protein (13±16% of the MG’s endogenous protein content), for which no ortholog could be found. The evolutionarily young gene coding this protein could be a promising candidate for antimicrobial research, with research into next-generation antibiotics often relying on antimicrobial peptides^20^. The gas-chromatography/mass-spectrometry (GC-MS) analyses of the MG further revealed 112 organic compounds (23 of which could not be identified, Extended Data Fig. 9, Supplemental Table 13). Six of the identified compounds had antibiotic- and/or fungicide-like structures and 35 were alkaloids. While we could not identify the exact structure of the alkaloids, many of them are known to have antimicrobial properties^21^. There were also 14 carboxylic acids, making up 52% of the secretions content (Extended Data Fig. 9, Supplemental Table 13), probably leading to a lower pH detrimental to bacterial survival and growth^6^.

The number of chemical compounds identified in the MG’s secretions of *M. analis* (112) is far greater than in other ant species, where the number of compounds ranges between 1 and 35 and mostly consists of carboxylic acids rather than alkaloids or antibiotic-like compounds^6^. While MG secretions have been observed to be used to sterilize the nest or as a response to fungal exposure in other ant species^6,8,10^, it had never been observed in the context of wound care. During wound care, the use of the MG secretions probably fulfils a similar role as the mammals’ antiseptic saliva. This analogous role likely led to the convergent evolution of functionally similar antimicrobial and wound healing proteins^22^. However, this treatment is indiscriminate in mammals as it has never been observed to depend on the infected state of the wound.

This study reveals a highly effective behavioral adaptation to identify and treat festering infections of open wounds in a social insect. The prophylactic and therapeutic use of antimicrobial secretions to counteract infection in *M. analis* (Fig. 3c) mirrors modern medical procedures used for treating dirty wounds^23^. Remarkably, the primary pathogen in ant’s wounds, *Pseudomonas aeruginosa*, is also a leading cause of infection in combat wounds, where infections can account for 45% of casualties in humans^24^. This demonstrates convergence in both the challenges of warfare and the solutions that evolved to mediate them in human and insect societies.

## Supporting information

Supplemental Model Workflow

## Methods

### Experimental design

The study was conducted in a humid savannah woodland located in the Comoé National Park, northern Côte d’Ivoire (Ivory Coast), at the Comoé National Park Research Station (8°46’N, 3°47’W)^25^. Experiments, observations, and sample collections in the Comoé National Park were carried out from April to June 2018, February, April to June and September to October 2019, April 2020 and October 2022. *Megaponera analis* is found throughout sub-Saharan Africa from 25°S to 12°N^26^ and known to show monophasic allometry within its worker sizes^27^. We thus divided the workers into majors (head width > than 2.40 mm), minors (head width < 1.99 mm) and intermediates (head width 2.40 - 1.99 mm). All experiments were carried out on the minor caste, the individuals most frequently injured^3^. All field studies were conducted in accordance with local legislation and permission by the Office Ivoirien des Parcs et Réserves (OIPR).

### Laboratory colonies

Eleven colonies, including queen and brood (colony size 1083±258 ants), were excavated and placed in artificial nests in the field stations laboratory. PVC nests (30×20×10 cm) were connected to a 1×1m feeding arena. The ground surface was covered with soil from the surrounding area (up to a height of 2 cm). Colonies were fed by placing in the feeding arena *Macrotermes bellicosus* termites collected from the surrounding area. These termites were found by scouts and triggered raiding behavior. The laboratory windows were kept open to maintain a natural humidity, temperature, and day-night cycle (light regime).

### Survival of injured ants

To quantify the lethality of various types of pathogens, workers were injured by a sterile cut in the middle of the femur on the hind leg and had the fresh wound submerged for 2 seconds in a 10 μL phosphate buffered saline (PBS) solution with a known pathogen concentration (approx. 10^6^ bacteria). Afterwards the injured ants were placed inside a cylindrical glass container with a diameter of 3 cm and a height of 5 cm. Before the experiments, the glass containers were filled with 1 cm of surface soil and placed for 3 hours at 220 °C in an oven together with the forceps and scissors for sterilization. Nest-like humidity was created by moistening the soil with 1 mL of sterilized water (boiled for ten minutes) and covered with aluminum foil. The isolation experiments were conducted at 24°C in a sterile room. To test for mortality, the injured ant was checked upon once per hour for the next 36 hours, if no reaction was observed even after shaking the container the ant was classified as dead. To ensure replicability of the survival experiments a negative (sterile PBS solution) and positive control (using a sterile PBS solution with 0.1 optical density (OD) of *Pseudomonas aeruginosa* (PSE)) were always included during each survival experiment. The sample sizes were PBS: *n*=104; PSE 0.1: *n*=61; PSE 0.05: n=15 (Fig. 2a & Extended Data Fig. 3). For all sample sizes and statistics see Extended Data Fig. 2.

To test if wound care by nestmates reduces the mortality of infected ants, we first marked 10 ants per colony during a raid. 24 hours later, all marked ants were injured in the same way as in the isolation trial. Afterwards, the wound of the injured ant was either exposed to a sterile PBS solution (sterile), a sterile PBS solution containing 0.05 OD of *P. aeruginosa* or a sterile PBS solution containing 0.1 OD of surface soil pathogens (grown on agar plates). Isolation trials with the same treatments were always conducted in parallel. Sample size of in nest survival experiments: PBS: *n*=12; PSE 0.05: *n*=12; Soil *n*=18 (Fig. 1c & 2a). For all sample sizes and statistics see Extended Data Fig. 2.

To test the importance of the Metapleural gland (MG) secretions we divided three colonies (approx. 1000 ants) into two equally sized sub-colonies including brood. The queens were left with a small retinue of 20 workers in a separate container. For each colony we plugged the MG opening of all workers of one of the sub-colonies with acrylic color, while for the other sub-colony acrylic color was placed on the thorax of the workers. Afterwards the experimental procedure was identical to the wound care experiment described above. Eight individuals were removed from each sub-colony, marked, and wounded and the wound exposed to either a sterile PBS solution (*n*=4 per sub-colony) or a sterile PBS solution containing 0.05 OD of *P. aeruginosa* (*n*=4 per sub-colony). Sample sizes were thus in total: Sterile: *n*=12; Infected: *n*=12 for both treatments (with and without plugged MG opening; Fig. 4). In addition, to quantify any effects of the plugging of the MG opening on individuals, a parallel experiment with *n*=12 Sterile and *n*=12 Infected ants with or without plugged MG opening was run in isolation (Extended Data fig. 5 & Supplemental Table 6).

The pathogen concentration was measured by optical density (OD) using a portable Ultrospec 10 cell density meter (Biochrom) with sterile PBS as solvent. For the soil pathogens, we collected surface soil in the surrounding area of the nest and grew it over 36 hours on agar plates (Extended Data Fig. 1b). For *P. aeruginosa* we created cultures of the isolated strains (Extended Data fig. 1c) in the field lab from frozen samples kept in Tryptic Soy Broth (TSB) medium with 25% glycerol (stored and transported at -23°C). After replating the bacterial culture once on a fresh plate, we waited 16 hours before applying the pathogen on fresh wounds. For all experiments, we used trypticase soy agar (TSA) plates to culture the bacteria.

### Treatment of wounds by nestmates

To quantify the wound care behaviors inside the nest, we filmed the ants using a Panasonic HC-X1000 and analyzed the videos using VLC media player v.3.0.16 Vetinari (intel 64bit). The wound of the injured ant was either exposed to a sterile PBS solution (Sterile) or a sterile PBS solution containing 0.05 OD of *P. aeruginosa* (Infected).

All manipulated ants were placed in front of the nest entrance directly after a raid and the nest was filmed for the subsequent 24 hours. Only one trial was conducted per colony with a total of four injured ants per colony (two sterile and two infected), in a total of three colonies (i.e., *n*=6 infected ants and *n*=6 sterile ants, Fig. 3b, c). The observed wound care behaviors were classified into two categories: (1) wound care: a nestmate cleans the open wound with its mouthparts; (2) metapleural gland (MG) care: a nestmate collects MG secretions either from its own gland (Supplemental Movie 1) or from the injured ant (Supplemental Movie 2) in its mouth before caring for the wound. These behaviors were quantified for the first 24 hours and summarized in 10 min intervals.

### Sample collection protocol for chemical, genetic and microbial analyses

To quantify the cuticular hydrocarbon (CHC) profile, differential gene expression (Extended Data Fig. 6) and pathogen load in the thorax content (Fig. 1a & 2b), we used the same experimental design as used to quantify the survival of injured ants (see above). In total, 30 sterile and 30 infected ants (*P. aeruginosa* OD=0.05) were prepared, 12 sterile and 12 infected ants were kept in isolation, another 12 sterile and infected ants were placed inside the nest and 6 sterile and 6 infected ants were collected immediately after injury. Sterile or infected ants were then collected from their enclosures at either 2 or 11 hours after manipulation (*n*=6 per treatment). Only ants that did not die until these time points were used for further analyses. The collected ants were then first placed in hexane for 10 minutes to extract the CHC profile. The gaster was then cut off and placed in RNAlater for genetic analyses and the thorax was placed in 100% ethanol for the microbial analyses. The samples for genetic and microbial analyses were brought to the University of Lausanne and the samples for chemical analyses to the University of Würzburg. Another 10 sterile and 10 infected ants kept in isolation were further observed for a total of 36 hours to ensure that the survival curves in this experiment resembled those in Fig. 2a.

### Pathogen identification & isolation

To isolate potential pathogens, we collected surface soil in the surrounding area of the nest and grew its water extract on Sarborough Dextrose Agar (SDA) plates for 36 h at local temperature to get the ‘soil pathogen mix’ (Extended Data Fig. 1b). Most of the plate was at first overgrown by black mycelia identified as *Rhizopus microsporus.* This fungus is known to contain symbiotic bacteria *Burkholderia* sp. (species not identified), as we confirmed by Sanger sequencing and bacterial community analysis (Fig. 1b). After several days, the culture started to show several large colonies of slimy microorganisms that were isolated and identified as *Pseudomonas aeruginosa* (Extended Data Fig. 1c). For infection assays (Extended Data Fig. 3), we isolated *Burkholderia* and its fungal host *Rhizopus* from each other by repeated passaging with antifungal nystatin (0.25 μl/ml in M9 agar, 30 °C) or antibacterial ciprofloxamin (0.02 mg/ml in SDA). No representatives of *Klebsiella* were isolated.

The identification of all microorganisms was done by preparation of a PCR-ready DNA from a piece of biomass as described in Lõoke et al.^28^. PCR reactions were done with universal fungal primers ITS5 5’-TCCTCCGCTTATTGATATGC-3’ and ITS4 5’-GGAAGTAAAAGTCGTAACAAGG-3’^29^ (GoTaq polymerase, Tm = 55 °C, elongation for 40 s) and commonly used universal bacterial primers 27F 5’-AGRGTTYGATYMTGGCTCAG-3’ and 1492R 5’-GGTTACCTTGTTACGACTT-3’ (same protocol but elongation for 1 min 20 s). The presence of PCR amplicons was verified on electrophoresis gel and the fragments were sent for Sanger sequencing. Resulting sequences were then screened against NCBI database with nucleotide BLAST^30^.

### Microbiome analysis and bacterial load quantification

To quantify the composition of the bacterial community in workers, we used the Powersoil DNA isolation kit (MO Bio) to extract DNA from *M. analis* thoraxes kept in RNAlater. Samples were homogenized by vortexing for 10 min, followed by two times 45 sec bead beating at 6 m/s using a FastPrep24^TM^ 5G homogenizer. Then we continued according to the kit’s manual and eluted DNA in 100 µL of nuclease-free water.

Bacterial loads were quantified with a QuantStudio5 qPCR instrument (Applied Biosystems) using the reaction set up described in Kešnerová et al.^31^ and the thermal cycling conditions recommended for SYBR® Select Master Mix. For quantifying total bacterial loads, we used our designed primers #1047 5’-AGGATTAGATACCCTRGTAGTC-3’ and #1049 5’-CATSMTCCACCRCTTGTGC-3’ (at doubled 0.4 µM concentration). For specific targeting we used *P. aeruginosa* primers #1209 5’-GTAGATATAGGAAGGAACACCAG-3’ and #1210 5’-GGTATCTAATCCTGTTTGCTCC-3’ and for normalization to the host’s housekeeping gene we used *M. analis* 28S rRNA gene primers #1207 5’-CTGCCCGGCGGTACTCG-3’ and #1208 5’-ACCGGGGACGGCGCTAG-3’. Serial dilutions (10x) showed that these primers performed with an amplification efficiency (E) of 1.86 (R^2^= 0.99), 2.00 (R^2^= 0.99), and 1.87 (R^2^= 0.99), respectively.

Total bacterial (Fig. 1a) and *P. aeruginosa* (Fig. 2b) 16S rRNA gene (target) copy numbers were expressed relatively to *M. analis* 28S rRNA gene copy numbers (host) based on the following equation: Δ Cq= 2^(Cqhost - Cqtarget), where Cq was the measured ‘quantification cycle’ value. To calculate the total 16S rRNA gene copy number in 1 μl of each DNA sample that was used for absolute bacterial abundance based on ASV counts (Fig. 1b) we used the equation n = E^(intercept – Cq)^32^, where standard curve’s *intercept* = 38.23^31^.

### 16S rRNA gene amplicon-sequencing

16S rRNA gene amplicon sequencing data were obtained from 40 experimental samples, a mock sample (to verify consistency of the MiSeq run compared to previous studies in our group), two blank DNA extractions and a negative PCR control with only H2O. We followed the Illumina 16S metagenomic sequencing preparation guide (https://support.illumina.com/documents/documentation/chemistry_documentation/16s/16s-metagenomic-library-prep-guide-15044223-b.pdf) to amplify and sequence the V4 region of the 16S rRNA gene. Primers for the first PCR step were 515F-Nex (TCGTCGGCAGCGTCAGATGTGTATAAGAGACAGGTGCCAGCMGCCGCGGTAA) and 806R-Nex (GTCTCGTGGGCTCGGAGATGTGTATAAGAGACAGGGACTACHVGGGTWTCTAA T). PCR amplifications were performed in a mix of 12.5 μL of Invitrogen Platinum SuperFi DNA Polymerase Master Mix, 5 μL of MilliQ water, 2.5 μL of each primer (5 μM), and 2.5 μL of template DNA. PCR conditions were 98 °C for 30 s, 25 cycles of 98 °C for 10 s, 55 °C for 20 s, and 72 °C for 20 s, and a final extension step at 72 °C for 5 min. Amplifications were confirmed by 2% agarose gel electrophoresis. The PCR products were then purified with Clean NGS purification beads (CleanNA) in a 1:0.8 ratio of PCR product to beads, and eluted in 27.5 μL of 10 mM Tris, pH 8.5. We then performed a second PCR step in which unique dual-index combinations were appended to the amplicons using the Nextera XT index kit (Illumina). Second-step PCRs were performed in a 25 μL, using 2.5 μL of the PCR products, 12.5 μL of Invitrogen Platinum SuperFi DNA Polymerase Master Mix, 5 μL of MilliQ water, and 2.5 μL of each of the Nextera XT indexing primers 1 and 2. PCR conditions were 95 °C for 3 min followed by eight cycles of 30 s at 95 °C, 30 s at 55 °C, and 30 s at 72 °C, and a final extension step at 72 °C for 5 min. The libraries were again purified using Clean NGS purification beads in a 1:1.12 ratio of PCR product to beads, and eluted in 27.5 μL of 10 mM Tris, pH 8.5. The amplicon concentrations, including the negative PCR control, mock and blanks, were quantified by PicoGreen and pooled in equimolar concentrations (except for the water and blank samples which were kept at lower concentrations). We verified that the final pool was of the right size using a Fragment Analyzer (Advanced Analytical). MiSeq (Illumina) sequencing was then performed at the Genomic Technology Facility of the University of Lausanne, producing (2 × 250 bp) reads. We obtained a total of 979’126 paired-end reads across the 40 experimental samples.

### Analyses of 16 rRNA gene amplicon-sequencing data

Raw sequencing data (deposited at the Sequence Read Archive (SRA) under PRJNA826317) were analyzed with the Divisive Amplicon Denoising Algorithm 2 (DADA2) package v.1.20.0 in R. All functions were run using the recommended parameters (https://benjjneb.github.io/dada2/tutorial.html) except for the filtering step in which we truncated the forward and reverse reads after 232 and 231 bp, respectively. At the learnErrors step, we then set randomize=TRUE and nbases=3e8. Amplicon-sequence variants (ASVs) were classified with the SILVA database (version 138). Unclassified ASVs and any ASV classified as chloroplast, mitochondria or Eukaryota were removed with the “phyloseq” package version 1.36.0, using the “subset taxa” function. We then used the “prevalence” method in the R package “decontam” v.1.12.0 to identify and remove contaminants introduced during laboratory procedures, using the negative PCR control and the blank samples as reference. This procedure filtered out three contaminant ASVs. Four additional ASVs were removed as they clearly represented (low abundance) contaminants due to index swapping from samples of a different project that were sequenced in the same sequencing run. We then calculated absolute bacterial abundances of each ASV by multiplying the proportion of each ASV by the total 16S rRNA gene copy number of each sample as measured by qPCR. To assess differences in community structure between treatments we ran ADONIS tests after calculating Bray-Curtis dissimilarities with the absolute ASV abundance matrix and plotted ordinations based on these Bray-Curtis dissimilarities (Extended Data Fig. 1a).

To test for differences in absolute abundance of individual bacterial genera between sterile and infected ants (Extended Data Fig. 1b), we used a permutation approach (permutation t-test) as done in Kešnerová et al.^33^. To do this, we selected the ASVs that had at least 1% relative abundance across five samples (18 ASVs belonging to 11 genera). We then calculated copy numbers at the genus-level for each of the 40 samples. At each time-point (2 h and 11 h), we randomized the values of the calculated copy numbers for each genus 10,000 times and computed the *t* values for the tested effect for each randomized dataset. The *P* values corresponding to the effects were calculated as the proportion of 10,000 *t* values that were equal or higher than the observed one.

### Chemical analysis of cuticular hydrocarbons

To quantify differences in CHC profiles between infected and sterile ant workers, cuticular hydrocarbon extracts were evaporated to a volume of approximately 100 μL and 1 μL was analyzed by using a 6890 gas chromatograph (GC) coupled to a 5975 mass selective detector (MS) by Agilent Technologies (Waldbronn, Germany). The GC was equipped with a DB-5 capillary column (0.25 mm ID × 30 m; film thickness 0.25 μm, J & W Scientific, Folsom, Ca, USA). Helium was used as a carrier gas with a constant flow of 1 mL/min. A temperature program from 60 °C to 300 °C with 5 °C/min and finally 10 min at 300 °C was employed. Mass spectra were recorded in the EI mode with an ionization voltage of 70 eV and a source temperature of 230 °C. The software ChemStation v. F.01.03.2357 (Agilent Technologies, Waldbronn, Germany) for windows was used for data acquisition. Identification of the components was accomplished by comparison of library data (NIST 17) with mass spectral data of commercially purchased standards and diagnostic ions.

To compare the relative abundances of the different compound groups (Extended Data Fig. 6), all compounds were identified (Supplemental Table 8) and grouped either into Alkanes, Alkenes, Alkadienes or Methyl-branched alkanes for each individual.

### Antimicrobial assay

To assess the antimicrobial efficacy of MG extracts (Fig. 3d), we quantified the increase over time of the OD in a 96-well plate box, with the outer row filled with 70 μL of PBS. In total we did 3 replicates per sample with a total of 70 μL per sample. Negative control: 70 μL Luria-Bertani (LB) broth (*n*=6). Positive control: 68 μL LB broth + 2 μL *P. aeruginosa* (*n*=6). MG sample: 66 μL LB broth + 2 μL *P. aeruginosa* + 2 μL MG sample (*n*=9).

For the preparation of the pathogen sample, we plated *P. aeruginosa* from a frozen stock on TSA plates. After 24 hours the bacterial culture was replated. After 12 hours aliquots were created using the new bacterial culture in a flask containing 15 mL of freshly sterilized LB broth for an initial OD reading. Afterwards the flask was placed in an incubator shaker (Amerex Steadyshake 757) at 180 rpm, 30 °C for the bacteria to grow. Once the OD readings were between 02-0.5 OD (the exponential growth phase) the flask was put on ice until the experiments started.

For the preparation of the MG sample, we pooled 10 MGs in 1.5 mL Eppendorf-Cups. We then froze the samples in liquid nitrogen and crushed the sample material using sterile pellets. We then added 50 μL of PBS-Buffer, vortexed shortly before centrifuging the sample for 5 min at 3000 rcf at 4 °C. We then extracted 30 μL of the supernatant into a new cup and repeated the centrifugation process. Afterwards 20 μL of the supernatant was placed again in a new cup and stored at -20 °C. Preparation of the wells was conducted on ice at 4 °C to prevent bacterial growth and degradation of the MG samples. The antimicrobial assays were done in a microplate reader (Synergy H1 BioTek) at 30 °C with a 600 nm wavelength for 8 hours with a double orbital shaking step after each OD reading cycle (every 10 min) at the University of Lausanne.

To calculate the intrinsic growth rate of the microbial population (r) in Fig. 3d we used the package growthcurver (v. 0.3.1) with the statistical software R v4.1.0. “r” represents the growth rate that would occur if there were no restrictions imposed on total population size.

### Sample collection for metapleural gland extracts

Due to the difficulty of collecting adequate amounts of metapleural gland secretions from the gland’s atrium (it is a very sticky substance that adheres to the cuticle), we decided to remove the atrium of the gland together with the secretory cells entirely, using a microscalpel and microscissors. To avoid using a solvent we used a Thermodesorber unit coupled to a GC-MS (TD-GC-MS). For this we transported frozen ants to the University of Würzburg and did the extractions directly in the laboratory next to the TD-GC-MS. One metapleural gland from each of six worker was pooled per sample. As a control we further collected pieces of cuticle of similar size from the side of the thorax to identify any potential contaminations which might have occurred during dissections. In total 3 samples (of 6 individuals each) were analyzed (Extended Data Fig. 8, Supplemental Table 12).

### Chemical analysis of the metapleural gland

The MG samples were placed in a glass-wool-packed thermodesorption tube and placed in the thermodesorber unit (TDU; TD100-xr, Markes, Offenbach am Main, Germany). The thermodesorption tube was heated up to 260°C for 10 min. The desorbed components were transferred to the cold trap (5 °C) to focus the analytes using N2 flow in splitless mode. The cold trap was rapidly heated up to 310 °C at a rate of 60 °C per minute, held for 5 min and connected to the GC-MS (Agilent 7890B GC and 5977 MS, Agilent Technologies, Palo Alto, USA) via a heated transfer line (300 °C). The GC was equipped with an HP-5MS UI capillary column (0.25 mm ID × 30 m; film thickness 0.25 μm, J & W Scientific, Folsom, Ca, USA). Helium was the carrier gas using 1.2874 ml/min flow. The initial GC oven temperature was 40 °C for 1 min, then raised at a rate of 5°C per min until reaching 300 °C, where it was held for 3 minutes. The transfer line temperature between GC and MS was 300 °C. The mass spectrometer was operated in electron impact (EI) ionization mode, scanning m/z from 40 to 650, at 2.4 scans per second. Chemical compounds were identified using the same protocol as for the CHCs.

### Proteomic analysis and sample preparation of the metapleural gland

To characterize the proteins secreted by the metapleural gland, we analyzed the proteome of the atrium of the gland, the metapleural gland secretory cells and the hemolymph (Extended Data Fig. 7). To be considered a metapleural gland protein that could mediate antimicrobial activity, proteins had to be found in the gland’s atrium, and must have a higher abundance in the gland’s atrium than in the hemolymph. The samples for proteomic analysis were collected in April 2020 from workers of field colonies collected in the Comoé National Park. Three types of samples were collected: Secretory: dissected secretory cells without the atrium, (*n*=6 samples, each pooled from five dissections); Hemolymph: hemolymph was collected from the thorax by a glass microcapillary through a small wound (*n*=6 samples, each pooled over five individuals, yielding 4-5 μL); Atrium: due to the difficulty of collecting pure MG content we chose to first widen the opening of the atrium and add 1 μl of PBS before extracting the content together with the PBS (*n*=6 samples, each pooled from five individuals). The samples were kept at -23 °C in Lo-bind Eppendorf tubes with 5μl of PBS together with a 1× Protease inhibitor Cocktail (Sigmafast) during transportation. Once in Lausanne, Switzerland, the samples were kept at -80 °C and processed swiftly.

Proteins were digested according to a modified version of the iST protocol^34^. Samples were resuspended in 20 μL of modified iST buffer (1% sodium deoxycholate, 10 mM DTT, 100 mM Tris pH 8.6) and heated at 95 °C for 5 min. They were then diluted with 24 μL of 4 mM MgCl2 and 1:100 of benzonase in H2O and incubated 15 min at ambient temperature. 14 μL of 160 mM chloroacetamide (in 10 mM Tris pH 8.6) were then added and cysteines were alkylated for 45 minutes at 25°C in the dark. After addition of 0.5 M EDTA (3 mM final concentration), samples were digested with 0.1 μg of trypsin/Lys-C mix (Promega) at 37 °C for 1 hour, followed by a second enzyme addition (0.1 μg trypsin/LysC) and 1h incubation. Two volumes of isopropanol + 1% trifluoroacetic acid (TFA) were added to one volume of sample and loaded onto an equilibrated OASIS MCX uElution plate (Waters) prefilled with SCX0 buffer (20% MeCN, 0.5% formic acid) and centrifuged. The columns were washed three times with 200 μL isopropanol + 1% TFA and once with 200 μL HPLC solvent A (2% MeCN, 0.1% formic acid). The peptide mixture was then sequentially eluted with 150 μL SCX125 buffer (20% MeCN, 0.5% formic acid, 125 mM ammonium acetate), 150 μL SCX500 buffer (20% MeCN, 0.5% formic acid, 500 mM ammonium acetate), and lastly with 150 μL basic elution buffer (80% MeCN, 19% water, 1% NH3).

Tryptic peptides fractions were dried and resuspended in 0.05% trifluoroacetic acid, 2% (v/v) acetonitrile, for mass spectrometry analyses. Tryptic peptide mixtures were injected on an Ultimate RSLC 3000 nanoHPLC system (Dionex, Sunnyvale, CA, USA) interfaced to an Orbitrap Fusion Tribrid mass spectrometer (Thermo Scientific, Bremen, Germany). Peptides were loaded onto a trapping microcolumn Acclaim PepMap100 C18 (20 mm × 100 μm ID, 5 μm, 100 Å, Thermo Scientific) before separation on a reversed-phase custom packed nanocolumn (75 μm ID × 40 cm, 1.8 μm particles, Reprosil Pur, Dr. Maisch). A flow rate of 0.25 μL/min was used with a gradient from 4 to 76% acetonitrile in 0.1% formic acid (total time: 65 min). Full survey scans were performed at a 120’000 resolution, and a top speed precursor selection strategy was applied to maximize acquisition of peptide tandem MS spectra with a maximum cycle time of 0.6 s. HCD fragmentation mode was used at a normalized collision energy of 32%, with a precursor isolation window of 1.6 m/z, and MS/MS spectra were acquired in the ion trap. Peptides selected for MS/MS were excluded from further fragmentation during 60 s.

Tandem MS data were processed by the MaxQuant software (version 1.6.14.0)^35^ using the Andromeda search engine^36^ matching to a custom-made protein database containing 19’618 sequences of *M. analis* (July 2019 version, see section “Genome annotation” for details), supplemented with sequences of common contaminants was used. Trypsin (cleavage at K,R) was used as the enzyme definition, allowing 2 missed cleavages. Carbamidomethylation of cysteine was specified as a fixed modification. N-terminal acetylation of protein and oxidation of methionine were specified as variable modifications. All identifications were filtered at 1% FDR at both the peptide and protein levels with default MaxQuant parameters. For protein quantitation the iBAQ values^37^ were used. MaxQuant data were further processed with Perseus software (version 1.6.14.0)^38^ for filtering, log2-transformation and normalization of values and ortholog annotations. To determine which proteins are secreted by the MG, we proceeded through a series of filters. Proteins not found in the atrium samples were eliminated and those found only in the MG atrium were considered hits. The log2 fold change between the atrium and the hemolymph were determined for all remaining proteins. Only proteins that were on average 1.5-fold more abundant in atrium samples than hemolymph were retained as hits. Percent of total iBAQ per sample was assigned to each protein to indicate abundance. Visualization of heatmaps was performed in Matlab 2020b (clustergram function). To determine whether our selected proteins were toxin-like, we ran clantox^39^ (http://www.clantox.cs.huji.ac.il/). Annotations (function, orthology depth and implications for wound healing) found in Supplemental Table 10 were determined using protein BLAST with the experimental clustered nr database^40^.

The mass spectrometry proteomics data are deposited at the ProteomeXchange Consortium via the PRIDE partner repository with the dataset identifier PXD033003.

### Genome sequencing and assembly

The genome of *M. analis* was sequenced and assembled by The Global Ant Genomics Alliance (GAGA, antgenomics.dk)^41^. To identify repeats in the genome assembly, we used the package RepeatModeler v2.0, which combines three *de-novo* repeat finding programs (RECON, RepeatScout and LtrHarvest/Ltr_retriever)^42^. RepeatModeler first probes chunks of the genome assembly to find repeats, then clusters and classifies identified repeats, producing a high-quality library of consensus sequences of repeated sequence families. The consensus sequences were used with RepeatMasker to softmask (in lower-case) all repeats in the genome assembly.

### Genome annotation

To annotate the *M. analis* genome assembly we used the *ab-initio* gene predictors Augustus v3.3.3^43^ and genemark^44^ within the BRAKER2 pipeline v2.1.4^45,46^. Both tools were trained using RNA-seq reads (see below) aligned to the softmasked version of the genome with STAR v2.7^47^ and protein evidence from NCBI refseq predictions for *Acromyrmex echinatior, Atta colombica, Camponotus floridanus, Dinoponera quadriceps, Linepithema humile, Monomorium pharaonis, Ooceraea biroi, Pogonomyrmex barbatus, Solenopsis invicta, Temnothorax curvispinosus, Vollenhovia emeryi and Wasmannia auropunctata*. To enable functional interpretation of experimental data, we annotated gene products. First, we identified gene orthologs in *Drosophila melanogaster* to leverage the excellent quality of the functional annotation in this species. To identify the orthologs, we applied a reciprocal best blast approach using Orthologr^48^ between the longest protein isoform for each gene from both species^49^. Additionally, we identified orthologs with *Apis mellifera* and *Camponotus floridanus* NCBI RefSeq gene annotations. Finally, we ran InterProScan (version 5.30-69.0 and panther-data-14.1) on all proteins using default settings. To assign functional information to *M. analis* annotated genes, we primarily used the gene ontology of *D. melanogaster* orthologs, but also results from sequence homology search on Interproscan and the Uniprot database.

To study the effect of infections on gene expression (Extended Data Fig. 7), we conducted a transcriptomic analysis. We dissected the gaster from the same infected (soil OD=0.1, *n*=20) and sterile ants (*n*=20), whose thorax were used for the microbiome analyses in Fig. 1AB. Samples were collected 2 and 11 hours after manipulation (*n*=10 per timepoint) in the Comoé field research station and had the sting together with the venom reservoir removed. The gaster was dissected without solvent and stored in a RNAlater stabilisation solution at -23 °C (ThermoFisher Scientific) for later analyses at the University of Lausanne. Afterwards, the samples were homogenized with ceramic beads in 1ml of Trizol reagent. Homogenized samples were incubated for 5 min at room temperature (RT) in Trizol, before adding Chloroform (200 μL). Samples were incubated for 5 min at RT then centrifuged (25 s at 12,000 rpm and 4 °C) and the upper aqueous layer (∼500 μL) transferred to a new tube. We added Isopropanol (650 μL) and Glycogen blue (1 μL; RNAse-free, Invitrogen, 15 mg/mL, #AM9516) then vortexed and incubated overnight at -20 °C. To purify the RNA, we used a EtoH precipitation method: samples were centrifuged (30 s at full speed at 4 °C), the supernatant was discarded and EtOH (1 mL at 80%) added. We repeated this step a second time with EtOH (1 mL at 70%). Finally, the supernatant was removed and the pellet, after a brief air dry (15-20 s) at RT, was resuspended in nuclease-free water. Libraries were prepared with the KAPA Stranded mRNASeq Library Preparation Kit (#KK8421) according to the manufacturer’s protocol. Paired end sequencing was performed on an Illumina Hiseq4000 sequencer at the Genomic Technology Facility of the University of Lausanne. We obtained ∼15-25 million PE reads per individual. Sequence reads have been deposited in NCBI Sequence Read Archive (SRA) under the accession number PRJNA823913.

We mapped RNA-seq reads on the softmask genome assembly using STAR v2.7 with the 2-pass mapping option and using the gene annotations to properly find exonic junctions^47^. Then, we used the program featureCounts from the Subread package to count the number of reads mapped on each exon for each gene^50^. Then using the count’s files, we performed differential gene expression analyses using DESeq2^51^ of sterile and infected workers collected at 2 and 11 hours after treatment. *P* values of differential expression analyses were corrected for multiple testing with a false discover rate (FDR) of 5%.

### X-Ray Micro-CT imaging

To examine the internal morphology of the MG (Fig. 3a), we scanned and visualized an intermediate sized worker (CASENT0744096), although multiple other workers of different sizes were scanned and visually inspected to confirm the morphology of the exemplar specimen was representative. The specimens were fixed in 90% ethanol, stained in 2 M iodine solution for a minimum of 7 days, then washed and sealed in a pipette tip with 99% ethanol for scanning. The scans were performed with the ZEISS Xradia 510 Versa 3D X-ray microscope at the Okinawa Institute of Science and Technology Graduate University, Japan. We scanned the thorax (mesosoma) and whole body to visualize both the structure of the gland, and its location in the whole ant. The scan parameters for the thorax scan were: Voxel size, 4.404 μm; Exposure time, 6 s; Voltage, 40 kV; Power, 3 W; Source distance, 23.147 mm; Objective, 4x; Projections, 1601. For the full body scan, we used: Voxel size, 12.5 μm; Exposure time, 1 s; Voltage, 40 kV; Power, 3 W; Source distance, 25.03 mm; Objective, 0.4x; Projections, 1601. The 3D reconstruction was performed using the ZEISS Scout-and-Scan Control System Reconstructor software (ZEISS Microscopy, Jena, Germany). The MG structures were manually segmented using Amira (version 2019.2; Thermo Fisher Scientific, Berlin, Germany) and visualized using VGStudio3.4 (Volume Graphics GmbH, Heidelberg, Germany). The gland atrium was rendered using isosurface and a clipping plane was created to visualize the cross-section. All other materials were visualized using Phong volume rendering.

### Statistical analysis

For statistical analyses and graphical illustration, we used the statistical software R v4.1.0^52^ with the user interface RStudio v1.4.1717 and the R package ggplot2 v3.3.5^53^. To select the appropriate statistical tests, we tested for deviations from the normal distribution with the Shapiro Wilks test (*P*>0.05). A Bartlett test was used to verify homoscedasticity (*P*>0.05). All our models included colony as a random factor and an overall likelihood ratio test against an intercept only model. In case of multiple testing, a Holm-Bonferroni correction was performed with the adjusted P-values given throughout the text. To test for significant differences in the survival curves, we conducted mixed effect cox proportional hazards regression models (Extended Data Fig. 2) using the R package survminer (v0.4.9) followed by post-hoc analyses using least square means with the R package lsmeans (v.2.30; Supplemental Table 2, 3 & 5). The survival curves were illustrated using Kaplan-Meier cumulative survival curves (Fig. 1c, 2a, 4, Extended Data Fig. 3 & 5). For the bacterial load (ι1Cq) within the thorax content between treated and untreated infected individuals (Fig. 1a & 2b) we conducted linear mixed effect models with a least square means (lsmeans) post-hoc analysis. For the bacterial growth inhibition (Fig. 3d) we conducted Mann-a Whitney U test. Differences in the CHC-profile composition were calculated using a permutational multivariate analysis of variance (ADONIS) on a Bray-Curtis dissimilarity matrix using the package vegan (v.2.5-7; Supplemental Table 7). For the different durations of MG care between sterile and infected individuals (Extended Data Fig. 4), we conducted a linear mixed effect model with individual as random factor followed by a Satterthwaite’s t-test. For differences between chemical compound groups of the CHC profiles (Extended Data Fig. 5), we conducted an analysis of variance (AOV) followed by a Tukey Honest Significant differences test (Supplemental Table 9). For behavioral differences in wound care between sterile and infected individuals (Fig 3b, c), we modeled wound care as a binary event using binomial generalized additive models with posthoc contrasts to identify intervals of time during which the probability of receiving wound care differed between sterile and infected individuals. See Supplemental Information for a reproducible modeling workflow.

### Data and materials availability

Raw amplicon-sequence data was deposited at the Sequence Read Archive (SRA) under PRJNA826317. Sequence reads were deposited in NCBI Sequence Read Archive (SRA) under the accession number PRJNA823913. The proteomics data was deposited at the ProteomeXchange Consortium via the PRIDE partner repository with the dataset identifier PXD033003. The remaining data is available in the Dryad Digital Repository: https://doi.org/10.5061/dryad.hqbzkh1j6

## End notes

### Acknowledgments

We thank the Comoé National Park Research Station for the use of their facilities for the field and laboratory research and the park management of Office Ivoirien des Parcs et Réserves for facilitating field research in the park. We thank GAGA for providing the assembled *Megaponera analis* genome. We thank Barbara Milutinovic and the Cremer lab at IST Austria for teaching us the methodology for the antimicrobial assay. We thank Christine La Mendola for her help with sample preparation for genetic analyses and Camille Lavoix for proof reading of the manuscript. This study was supported by the Swiss NSF and an ERC grant to LaK. ACL was supported by Swiss SNSF grant PR00P3_179776. ETF was also supported by the DFG Emmy Noether Programme (n°: 511474012).

### Author contributions

Conceptualization: ETF, LaK

Methodology: ETF, LuK, JL, QH, ACL, AD, FA, EPE, PW, TS, DBS

Investigation: ETF, LuK, JL, QH, ACL, EPE, TS

Visualization: ETF, QH, FA, EPE, ACL, JL, DBS

Funding acquisition: LaK

Project administration: ETF, PE, LaK Supervision: ETF, LaK

Writing – original draft: ETF, LaK

Writing – review & editing: ETF, LuK, JL, QH, ACL, AD, FA, EPE, PW, PE, TS, DBS, LaK

### Additional information

Supplementary information is available for this paper. Correspondence and requests for materials should be addressed to Erik T. Frank.

## Extended Data

**Extended Data Fig. 1.**
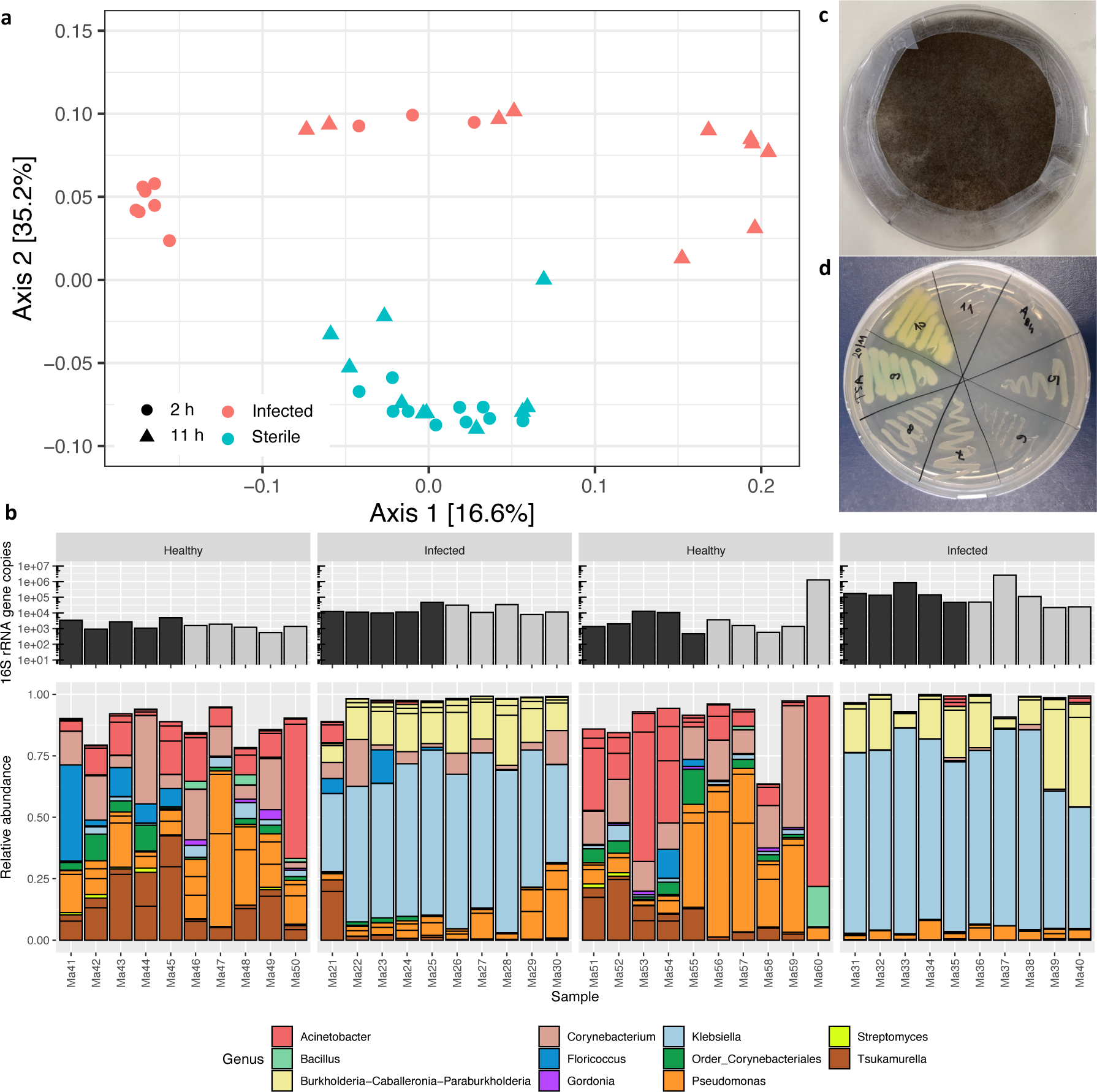
Ant microbiome and soil pathogens. **(a)** Principal Coordinate Analysis of the microbiome of the thorax of sterile and infected ants at 2 and 11 hours (as seen in Fig. 1b). ADONIS: Treatment: *F*=17.451; *R*^2^=0.31; *P*<0.001. **(b)** 16S rRNA gene copy numbers and relative abundance of bacterial genera present in the thorax of the same individuals as in Fig. 1A. Multiple bars of the same color indicate different amplicon-sequence variants (ASVs) belonging to the same genera. ADONIS: Treatment: *F*=17.45; *R*^2^=0.31; *P*<0.001 **(c)** Microbial culture of surface soil grown on an agar plate (mostly showing fungal mycelia of *Rhizopus* with black spores). **(d)** Bacterial cultures of isolated *Pseudomonas aeruginosa* strains from the soil.

**Extended Data Fig. 2.**
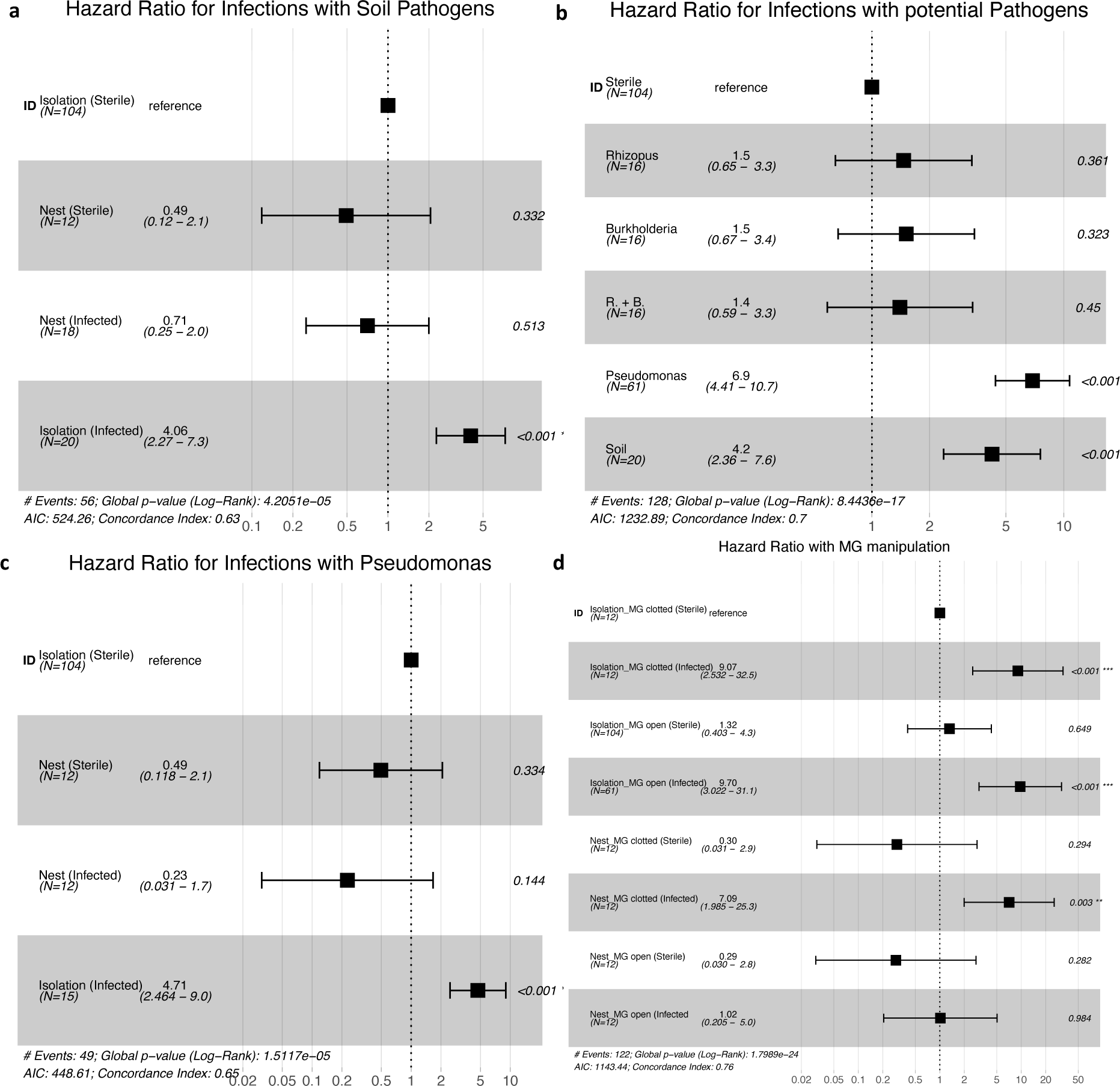
Cox proportional hazard models for survival of infections. The reference were sterile ants in isolation with colony of origin as random factor. **(a)** Statistical results for infected ants (surface soil pathogens: OD=0.1) in isolation and inside the nest (Fig. 1c). **(b)** Statistical results for infected ants kept in isolation exposed to different potential pathogens cultured from the soil (*Burkholderia, Rhizopus, Pseudomonas*; OD=0.1) or surface soil pathogens (Extended Data Fig. 3). **(c)** Statistical results for infected and sterile ants (*P. aeruginosa*: OD=0.05) in isolation and with nestmates in the nest (Fig. 2a). **(d)** Statistical results of infected ants (*P. aeruginosa*: OD=0.05) kept in isolation or inside sub-colonies containing workers with or without plugged MG openings (Fig. 4; Extended Data Fig. 5).

**Extended Data Fig. 3.**
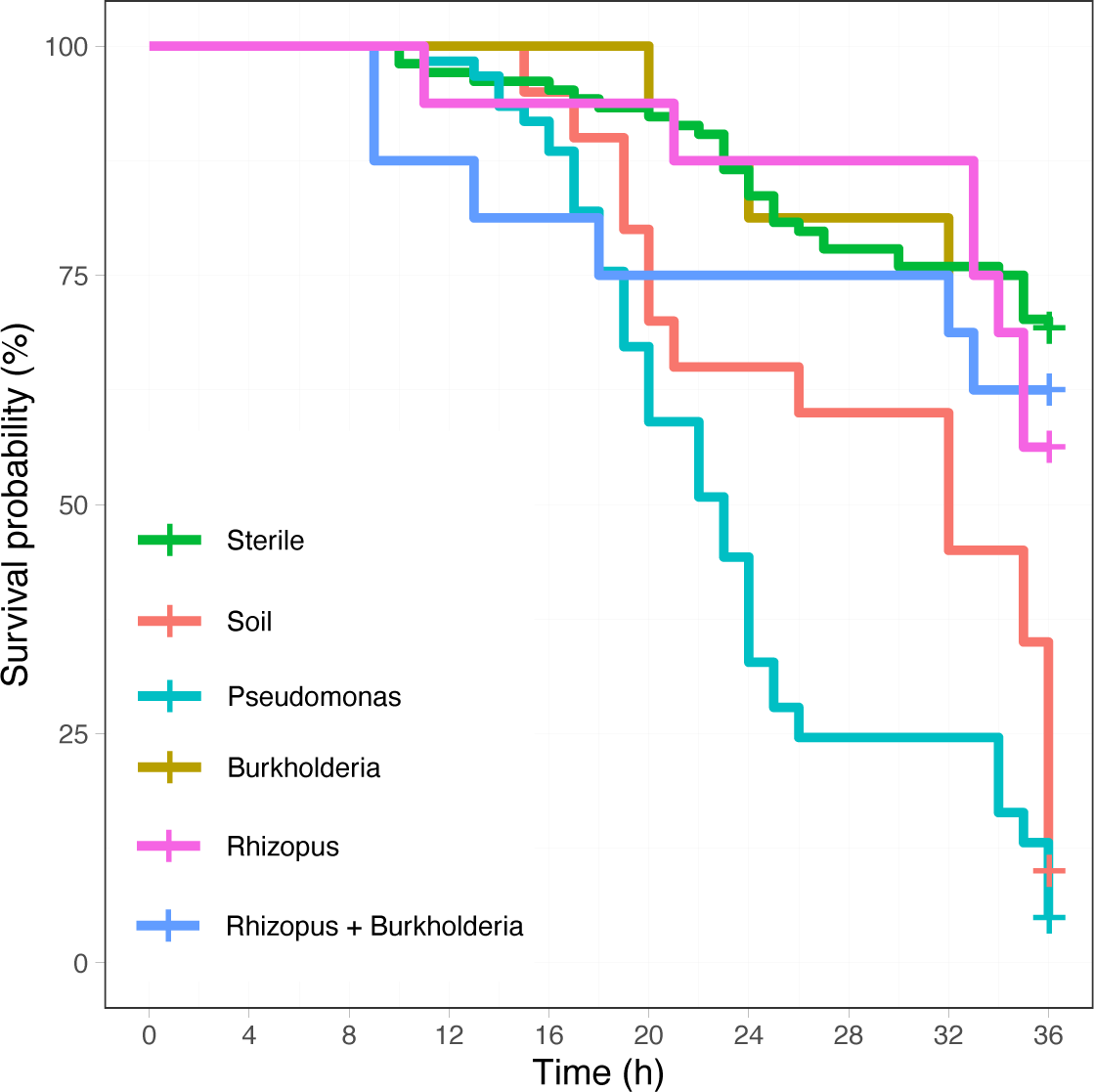
Survival probability for different pathogen types. Kaplan – Meier cumulative survival rates of workers kept in isolation chambers. All individuals had one hind leg cut off at the femur, the wound was then exposed to a sterile PBS solution (Sterile, *n*=104), a mix of either surface soil pathogens (Soil OD=0.5, *n*=20) or isolated pathogens diluted in PBS (OD=0.1): *P. aeruginosa* (*Pseudomonas, n*=61), *Burkholderia* (*n*=16), *Rhizopus* (*n*=16), or *Rhizopus* with *Burkholderia* (*n*=16). The treated ants were observed for 36 hours and the time of death noted. Statistical significance tested with a mixed-effects Cox proportional hazards regression model (Extended Data Fig. 2b).

**Extended Data Fig. 4.**
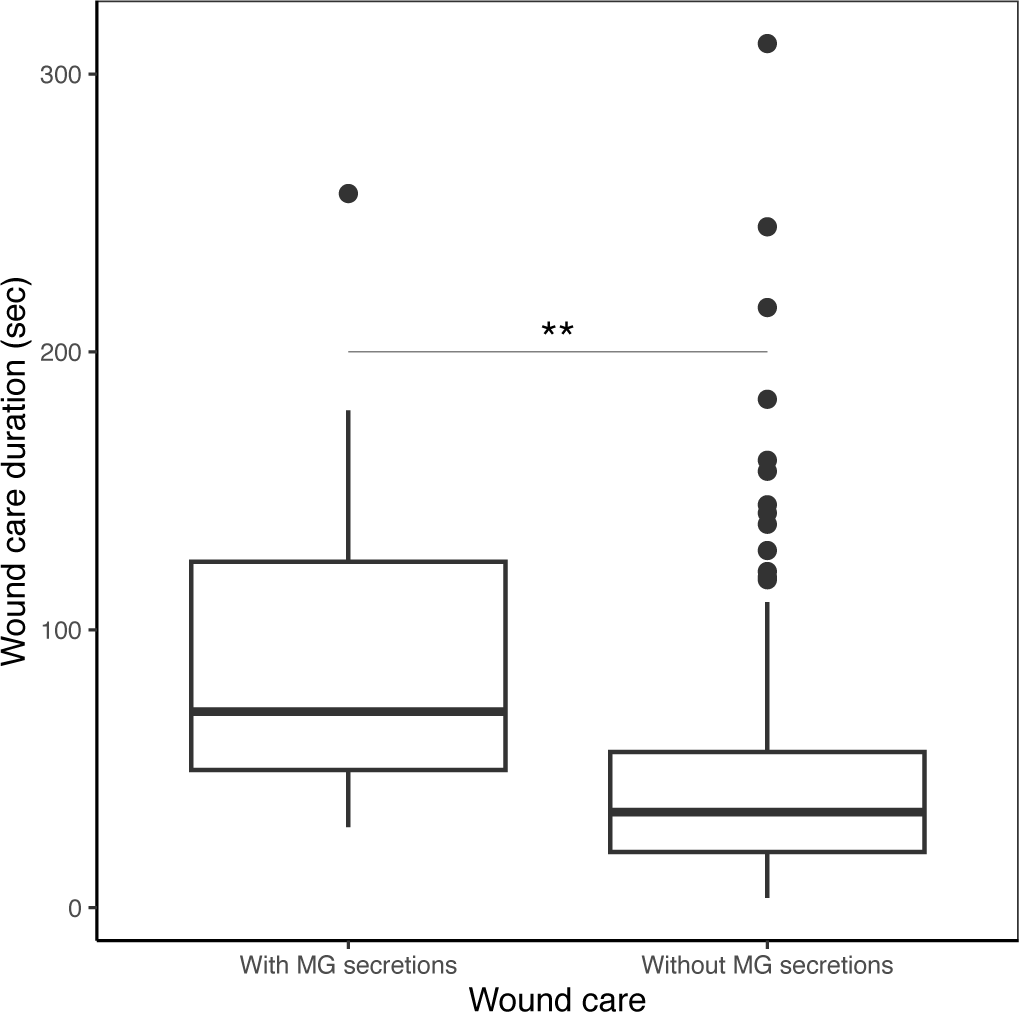
Differences in wound care duration with and without metapleural gland secretions. Duration of wound care with and without MG secretions for the first 24 hours after injury inside the nest. With MG secretions: wound care involving the MG for both infected and sterile ants pooled together using either the MG of the injured ant (*n*=26 for infected, *n*=10 for sterile ants) or of the nursing ant (*n*=2 for infected, *n*=4 for sterile ants), total sample size *n*=42. Without MG secretions: wound care excluding the MG for both infected and sterile ants pooled together (*n*=174 for infected, *n*=192 for sterile ants), total sample size *n*=366. Linear mixed effect model (Random Factor: Individual: Variance=134.4, Std. Dev.=11.59; Residual: Variance=2671.2, Std. Dev.=51.68) with Satterthwaite’s t-test for wound care duration with and withut MG secretions: t=-3.09; *P*=0.003**.

**Extended Data Fig. 5.**
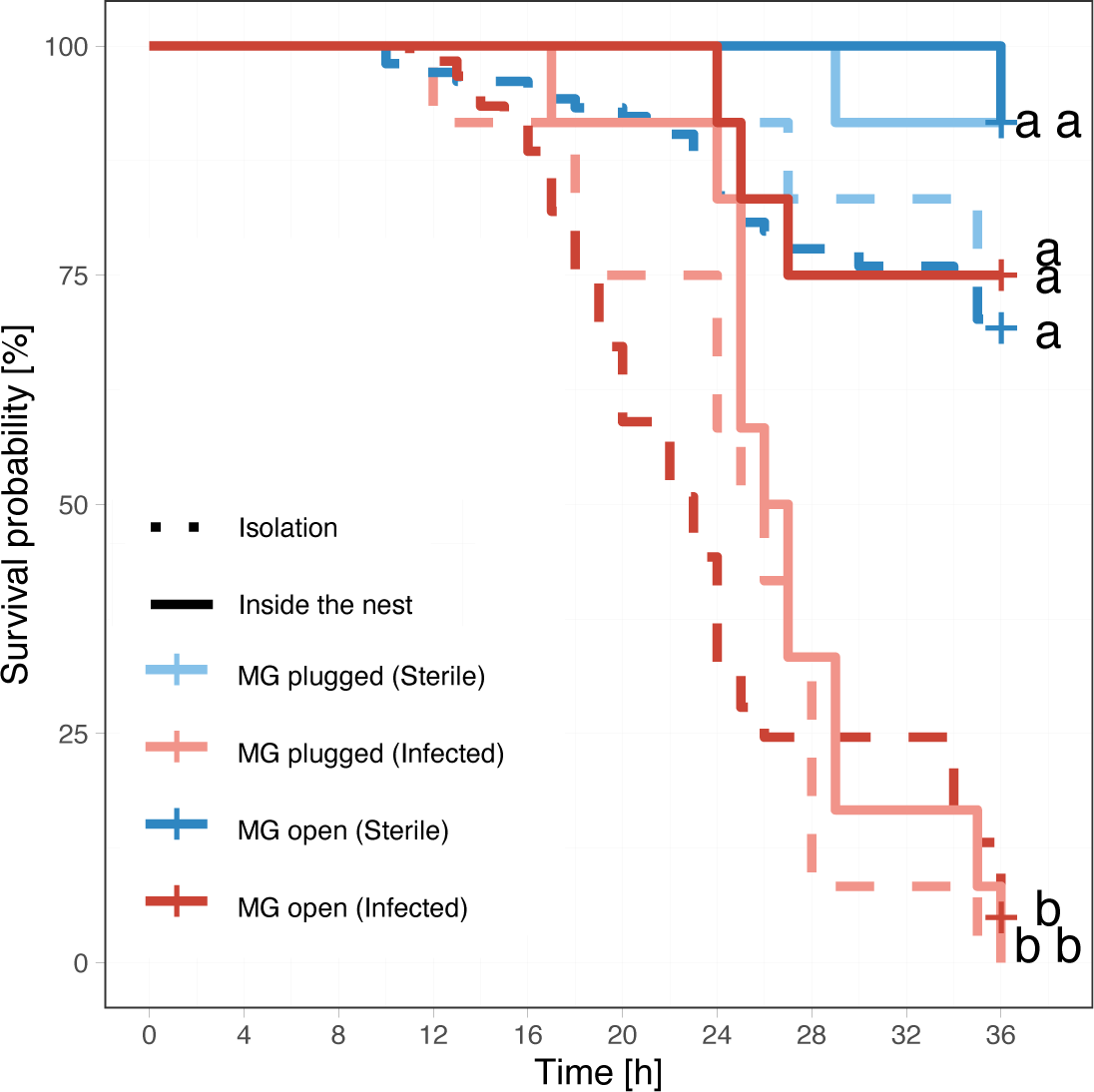
Importance of MG secretions on survival of sterile and infected ants. Kaplan – Meier cumulative survival rates of workers in isolation (dotted line) or inside sub-colonies (solid line) whose wounds were exposed to *P. aeruginosa* diluted in PBS (Infected, OD=0.05) or a sterile PBS solution (Sterile). In three sub-colonies all ants had their MG opening closed with acrylic color (MG plugged), while it remained open in the other three sub-colonies (MG open). *n*=12 per treatment. Detailed statistical results in Extended Data Fig. 2d and Supplemental Table 5.

**Extended Data Fig. 6.**
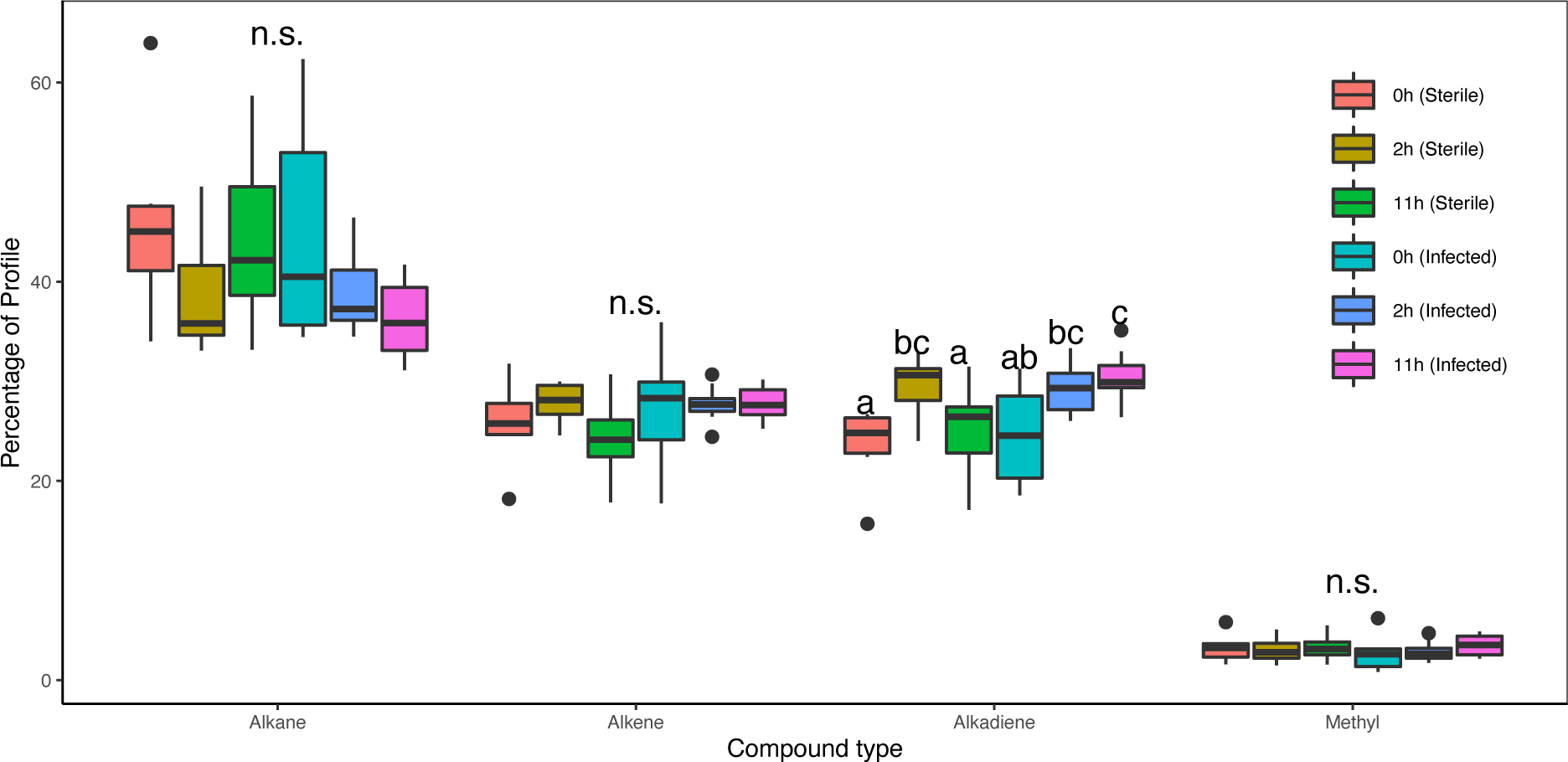
Chemical composition of cuticular hydrocarbon profiles of sterile and infected ants. Differences across compound types (alkanes, alkenes, alkadienes and methyl-branched alkanes) for sterile ants (Sterile; *n*=30) and ants infected with *P. aeruginosa* diluted in PBS (OD=0.05, Infected; *n*=30) at three different time points (0, 2 and 11 hours). Detailed statistical results in Supplemental Table 8.

**Extended Data Fig. 7.**
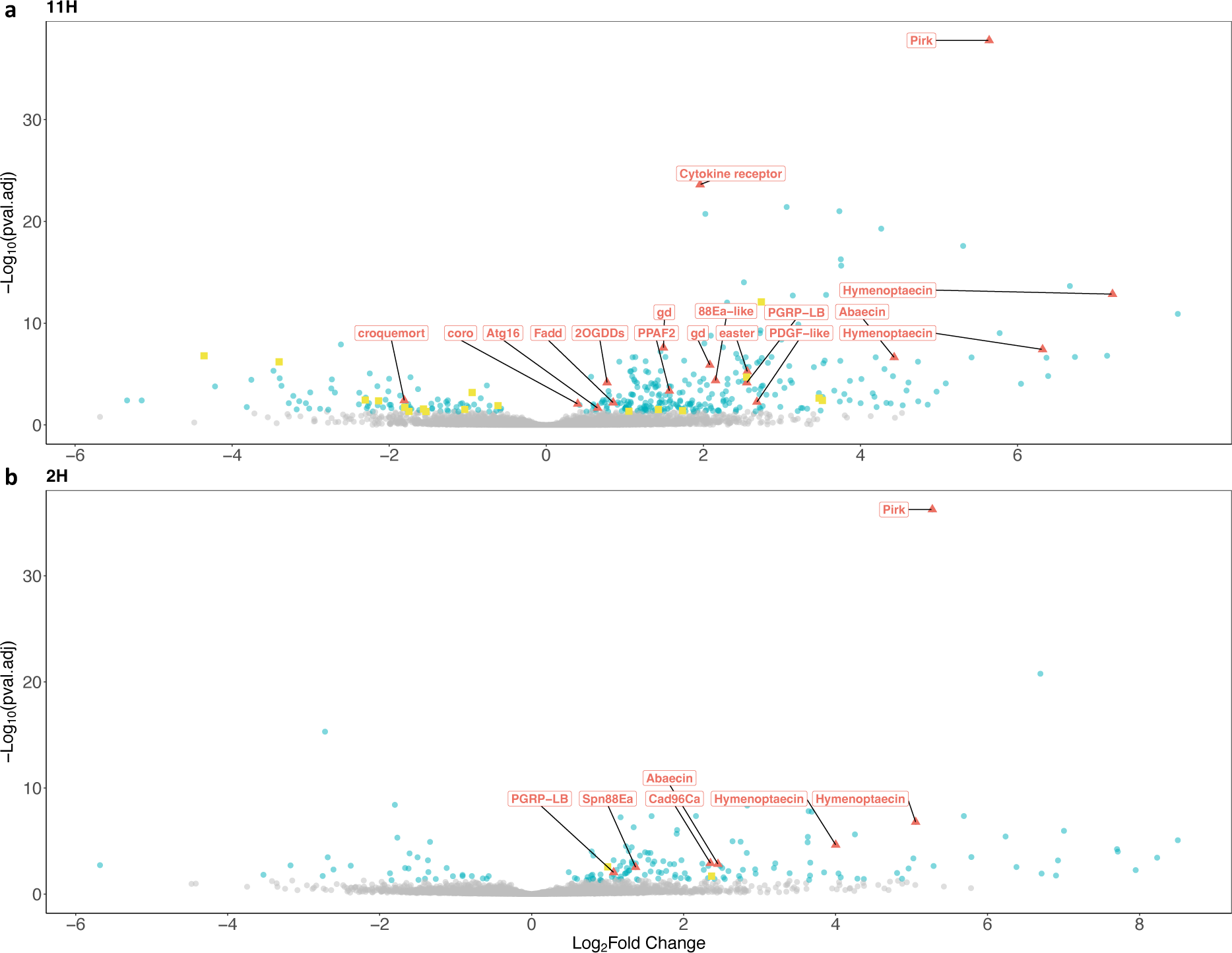
Differential gene expression between sterile and infected ants 2 and 11 hours after injury. Infection triggers changes in the expression of hundreds of genes. Volcano plot illustrates the fold up- and down-regulation of immune-related genes (red triangles, Supplemental Table 10) and lipids and CHC-related genes (yellow squares, Supplemental Table 9), 11 hours **(a)** and 2 hours **(b)** after infection. Positive Log2FoldChange values correspond to genes up-regulated in infected ants when compared to sterile ants, while negative values are down-regulated in infected ants. *P*-values are corrected for multiple testing using the Benjamini and Hochberg method (genes with significant differential expression marked in blue, i.e. adjusted *P*-value <0.05).

**Extended Data Fig. 8.**
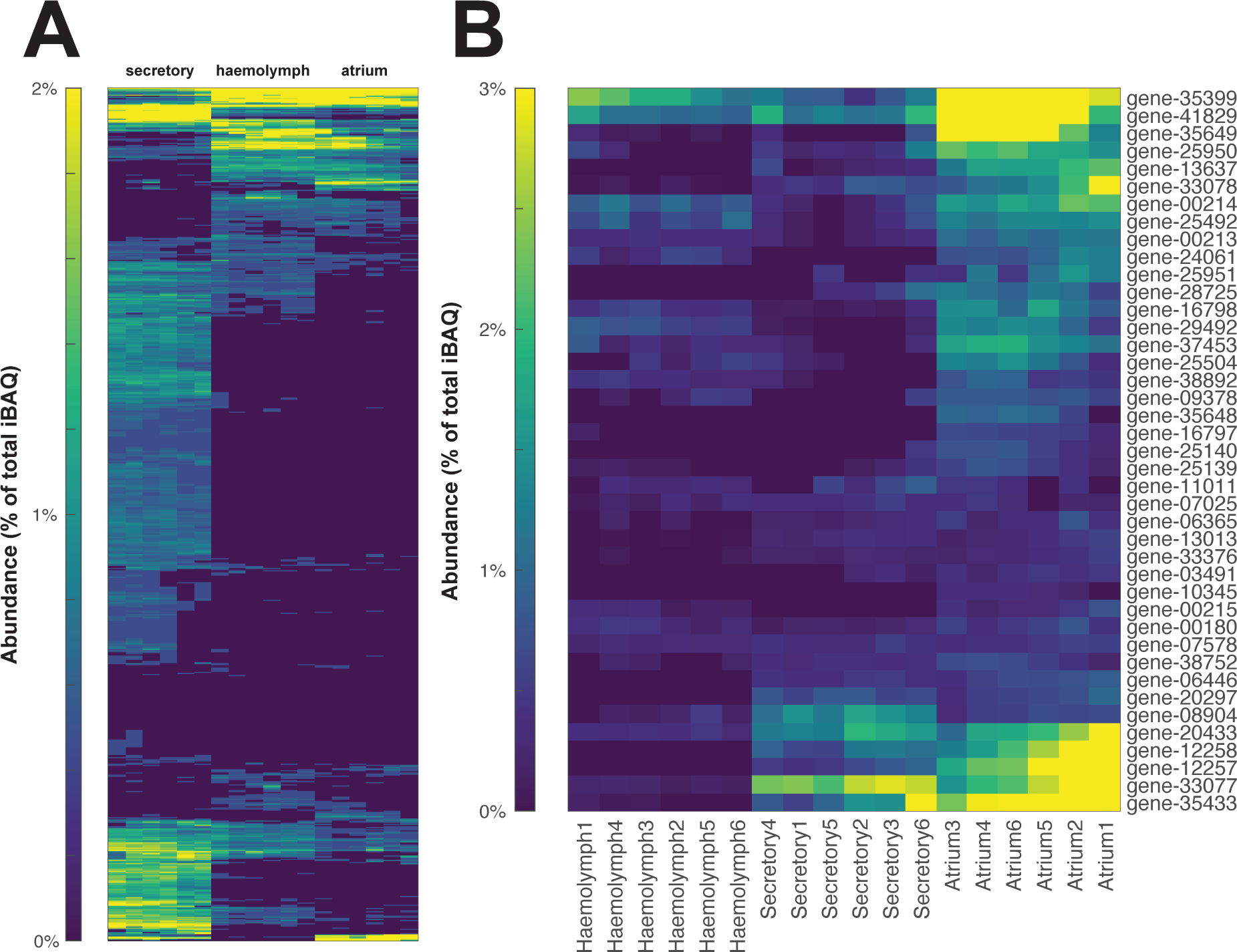
Proteins from the metapleural gland. **(a)** All proteins identified in hemolymph (*n*=6), secretory cells (*n*=6) and metapleural gland atria (*n*=6). **(b)** The 41 proteins deduced through filtering to be secreted by the metapleural gland. Gene annotations can be found in Supplemental Table 11. Heatmap colors indicate normalized percentages of total iBAQ for a given sample.

**Extended Data Fig. 9.**
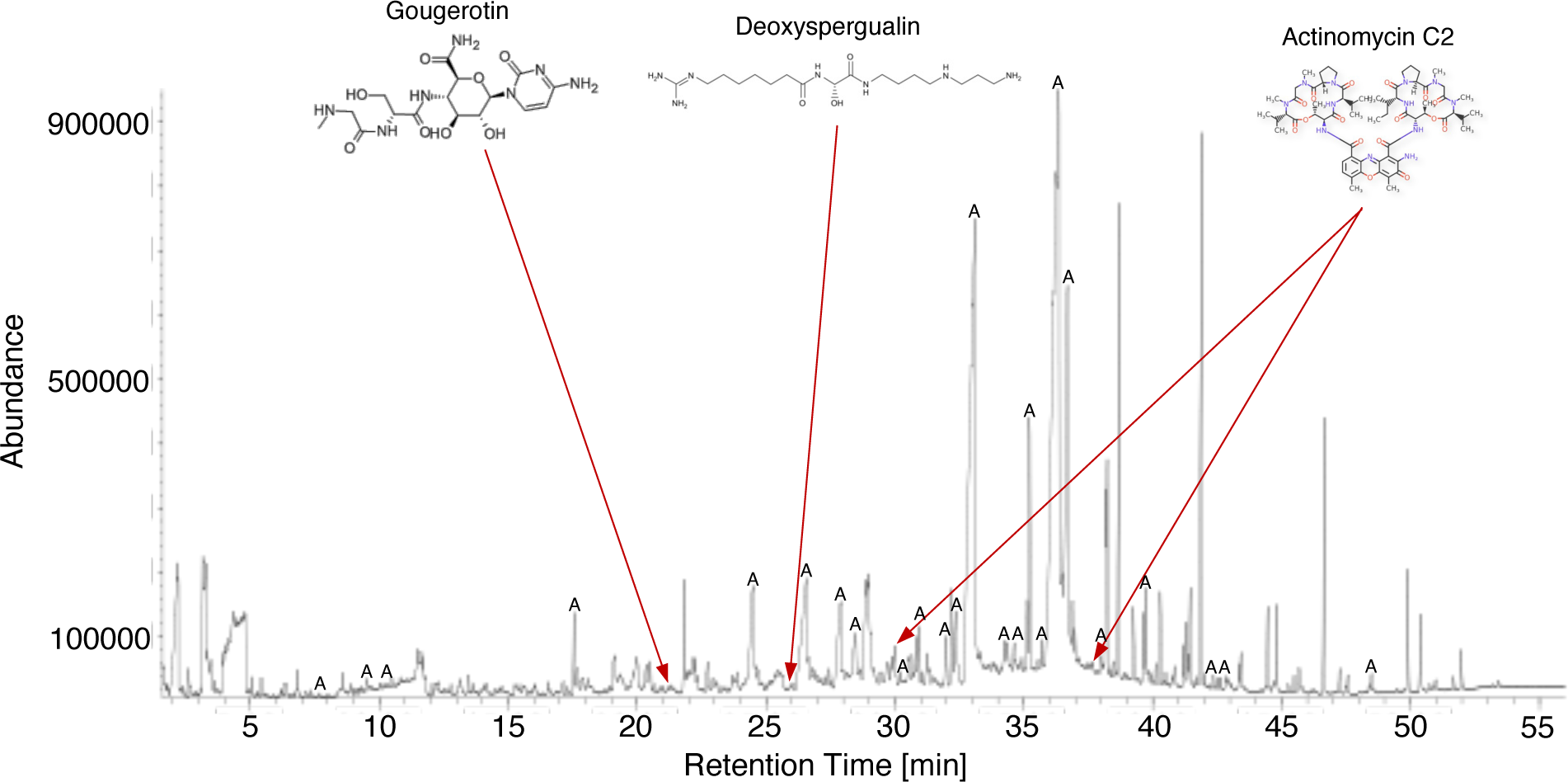
Chemical compounds in the metapleural gland. Gas-chromatographic representation of one sample of 6 pooled metapleural glands. Red arrows indicate chemical compounds with similar structure to known antibiotics and antimicrobials. Peaks marked with the letter A represent carboxylic acids. A detailed list of all chemical compounds found in the MG is described in Supplemental Table 12.

## Supplemental Materials

Supplemental Tables 1 to 13

Supplemental Model Workflow

Supplemental Movies 1 to 2

