## Supplemental Model Workflow for "Antimicrobial wound care in an ant society"

Appendix 1: Hierarchical generalized additive modeling

**References** **11**

### Summary

This appendix provides a reproducible workflow for our analysis of wound care using hierarchical generalized additive models (Pedersen et al. 2019). These models test the hypothesis that ants of the species *Megaponera analis* can perceive whether wounded nestmates are infected and direct their wound care accordingly. This hypothesis predicts that infected ants should receive more wound care than uninfected ants.

We find that infected ants were more likely to receive wound care than uninfected ants, both in terms of total wound care and wound care specifically involving metapleural gland (MPG) secretions, during a period between 10 and 12 hours after wounding. This supports the hypothesis *M. analis* can perceive whether a wounded nestmate is infected and use this information to decide whether to provide care, but it also suggests that there is a critical period between of time when such preferential care is given.

---

### Prepare environment

```
library(tidyverse)
library(lubridate)
library(mgcv)
library(gratia)
library(DHARMa)
library(patchwork)
library(emmeans)
library(marginaleffects)
```

### Load data

We will treat total wound care separately from the subset of wound care interactions involving MPG secretions.

#### Total wound care

```
total_wound_care <- read_csv("../data/total_wound_care.csv") %>%
  # rename columns
  select(date = Date, color.code = "Colour code", time = Time, colony = Colony,
         id = ID, trt = Treatment, t.allo = "t allo", allo = Allo,
         t.wound = "t wound", wound = Wound) %>%
  # process variables
  mutate(date = dmy(date),
         colony = factor(colony),
         id = factor(id),
         hours = time/60, # time in hours
         t.wound.bin = as.integer( # binary wound care
           if_else(t.wound > 0, 1, 0)
           ),
         trt = factor(if_else(trt == "PBS", "Sterile", trt))
         )
```

##### MPG wound care

```
mpg_wound_care <- read_csv("../data/mpg_wound_care.csv") %>%
  # rename columns
  select(colony = Colony, id = Individual, trt = Treatment,
         hours = Timeslot, mpg = MPG) %>%
  # process variables
  mutate(colony = factor(colony),
         id = factor(id),
         mpg.wound.bin = as.integer( # binary wound care
           if_else(mpg > 0, 1, 0)
           ),
         trt = factor(if_else(trt == "PBS", "Sterile", trt))
         )
```

##### Predictor matrix

This matrix of the predictor variables used in our models will later be passed to `predict()` to generate model predictions over the joint range of our predictor variables.

```
nd <- crossing(trt = c("Infected", "Sterile"),
               hours = seq(0, 24, 0.1),
               id = 0,
               colony = 0)
```

##### Inverse logit function

This will be used to back-transform the predictions of our binomial models, which are fit using a logit link function.

```
inv_logit <- function(x) {
  x <- 1/(1+(exp(-x)))
}
```

#### Modeling

We have a binary response variable measured across treatments and through time, and measurements are further grouped by colony and individual. What we want is the treatment * time interaction, but we need to deal appropriately with the colony/individual group structure. This calls for an HGAM approach capable of capturing nonlinearity in the temporal patterns while respecting the nested grouping of the measurements. These models follow the type I form described by Pedersen et al. (2019) in which independent smooths are estimated for each level of treatment. When models are specified in this way, treatment must also be included as a fixed intercept effect. Individual and colony are included as nested random intercept effects. Models are implemented using `mgcv` (Wood 2017).

##### Call models

```r
# All care
gam_00 <- gam(t.wound.bin ~
                # treatment as fixed intercept effect
                trt +
                # individual and colony treated as nested random effects
                s(id, colony, bs = "re") +
                # interaction smooth for time * treatment
                s(hours, by = trt, bs = "tp", k = 10),
              family = "binomial",
              method = "REML",
              data = total_wound_care)

# MPG care
gam_10 <- gam(mpg.wound.bin ~
                # treatment as fixed intercept effect
                trt +
                # individual and colony treated as nested random effects
                s(id, colony, bs = "re") +
                # interaction smooth for time * treatment
                s(hours, by = trt, bs = "tp", k = 10),
              family = "binomial",
              method = "REML",
              data = mpg_wound_care)
```

#### Validate models

We validate models using `DHARMa` (Hartig 2021). The total care model passes validation tests, and the MPG care model shows only weak deviations from distributional assumptions.

```r
# All care
gam_00_resid <- simulateResiduals(gam_00)
plot(gam_00_resid)
```

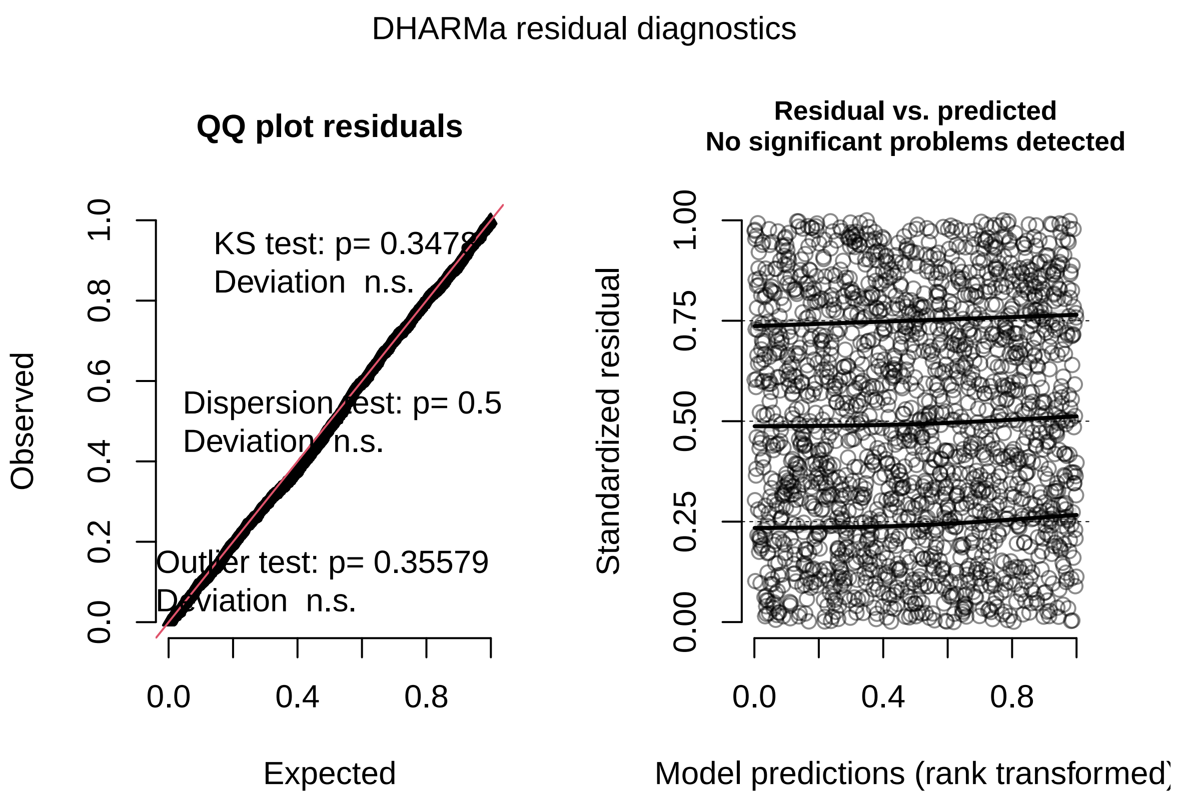

```
# MPG care
gam_10_resid <- simulateResiduals(gam_10)
plot(gam_10_resid)
```

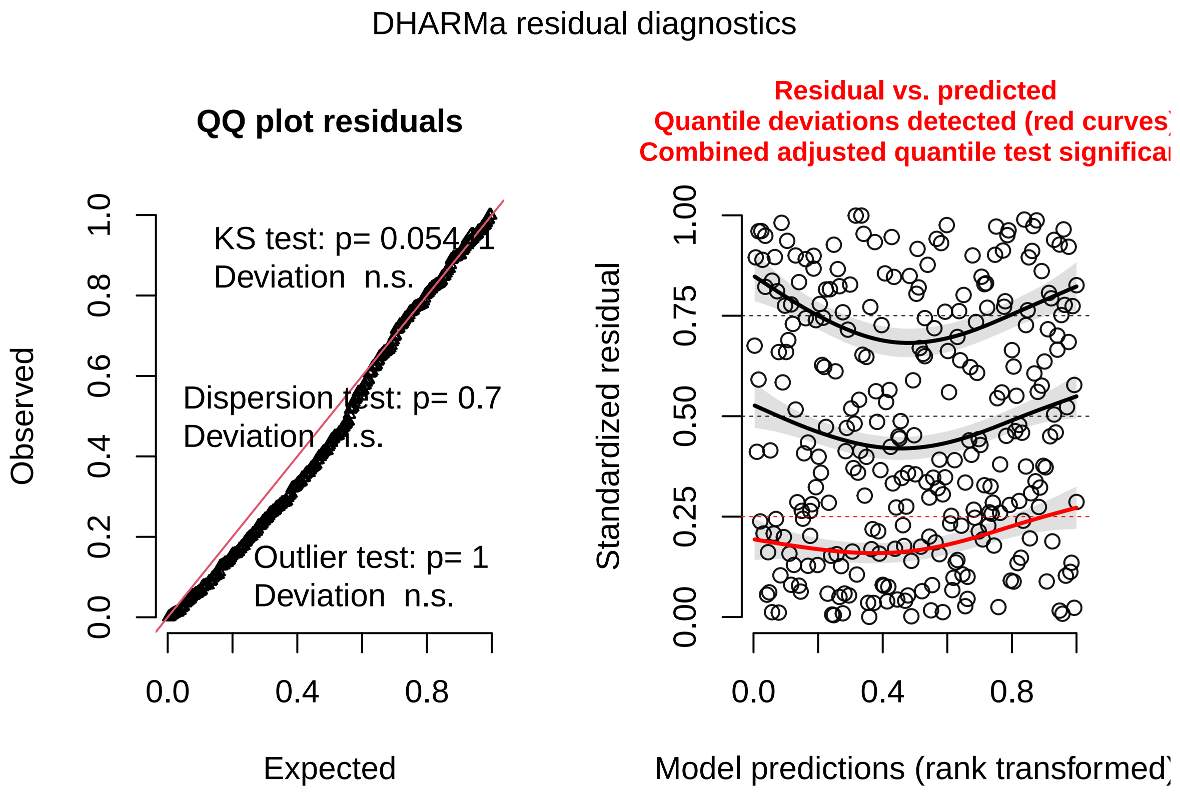

##### Tabulate output

Note that the "significance" reported for smooths refers to the null hypothesis that the true smooth function is a flat line. The test is performed on the `edf` estimate, which corresponds approximately to the degree of a polynomial function that might be fit to the data (e.g. `edf = 3` would indicate curve complexity comparable to that of a cubic polynomial function). This is not a test for differences in smooths between treatments. This must be tested separately via a smooth contrast, performed below.

We see strong nonlinearity in the binomial response for all care across both treatments. For MPG care, the curves are flatter, and only the curve for infected individuals is significantly different from a flat line.

The fixed intercept effect of treatment was not significant in either model, indicating that there is no difference between treatments in the global mean of the binomial response.

```
# All care
summary(gam_00)
```

```
##
## Family: binomial
## Link function: logit
##
## Formula:
## t.wound.bin ~ trt + s(id, colony, bs = "re") + s(hours, by = trt,
##     bs = "tp", k = 10)
##
```

```
## Parametric coefficients:
##              Estimate Std. Error z value Pr(>|z|)
## (Intercept) -1.500527   0.329048  -4.560 5.11e-06 ***
## trtSterile  -0.001127   0.463657  -0.002    0.998
## ---
## Signif. codes:  0 '***' 0.001 '**' 0.01 '*' 0.05 '.' 0.1 ' ' 1
##
## Approximate significance of smooth terms:
##                        edf Ref.df Chi.sq p-value
## s(id,colony)         9.211 10.000  76.18  <2e-16 ***
## s(hours):trtInfected 7.123  8.157  97.95  <2e-16 ***
## s(hours):trtSterile  4.581  5.616  66.32  <2e-16 ***
## ---
## Signif. codes:  0 '***' 0.001 '**' 0.01 '*' 0.05 '.' 0.1 ' ' 1
##
## R-sq.(adj) =  0.166   Deviance explained = 16.9%
## -REML = 822.39  Scale est. = 1         n = 1833
```

```
# MPG care
summary(gam_10)
```

```
##
## Family: binomial
## Link function: logit
##
## Formula:
## mpg.wound.bin ~ trt + s(id, colony, bs = "re") + s(hours, by = trt,
##     bs = "tp", k = 10)
##
## Parametric coefficients:
##             Estimate Std. Error z value Pr(>|z|)
## (Intercept) -2.39984    0.46195  -5.195 2.05e-07 ***
## trtSterile  -0.08747    0.58844  -0.149    0.882
## ---
## Signif. codes:  0 '***' 0.001 '**' 0.01 '*' 0.05 '.' 0.1 ' ' 1
##
## Approximate significance of smooth terms:
##                        edf Ref.df Chi.sq p-value
## s(id,colony)         2.197 10.000  2.726 0.25857
## s(hours):trtInfected 4.803  5.850 18.314 0.00409 **
## s(hours):trtSterile  1.543  1.903  4.727 0.05968 .
## ---
## Signif. codes:  0 '***' 0.001 '**' 0.01 '*' 0.05 '.' 0.1 ' ' 1
##
## R-sq.(adj) =  0.155   Deviance explained = 21.2%
## -REML =  93.04  Scale est. = 1         n = 288
```

#### Calculate contrasts

We will use `marginaleffects` (Arel-Bundock 2023) to identify portions of the timer series during which the probability of receiving care differs between infected and sterile ants.

```r
# Total care
gam_00.contrast <- avg_comparisons(gam_00,
                                   variables = c("trt"),
                                   newdata = nd,
                                   by = "hours",
                                   type = "response")
```

```
## Warning: The `trt` variable is treated as a categorical (factor) variable, but
##   the original data is of class character. It is safer and faster to
##   convert such variables to factor before fitting the model and calling
##   `slopes` functions.
##
##   This warning appears once per session.
```

```r
gam_00_contrast_intervals <- gam_00.contrast %>%
  mutate(sig = if_else(conf.low > 0 | conf.high < 0, TRUE, FALSE)) %>%
  select(hours, sig) %>%
  mutate(index = row_number()) %>%
  mutate(flag = if_else(lag(sig) != sig, 1, 0) %>% coalesce(0)) %>%
  mutate(flag_cum = factor(cumsum(flag))) %>%
  filter(sig == TRUE) %>%
  group_by(flag_cum) %>%
  summarize(start = min(hours), end = max(hours)) %>%
  select(-flag_cum)

# MPG care
gam_10.contrast <- avg_comparisons(gam_10,
                                   variables = c("trt"),
                                   newdata = nd,
                                   by = "hours",
                                   type = "response")

gam_10_contrast_intervals <- gam_10.contrast %>%
  mutate(sig = if_else(conf.low > 0 | conf.high < 0, TRUE, FALSE)) %>%
  select(hours, sig) %>%
  mutate(index = row_number()) %>%
  mutate(flag = if_else(lag(sig) != sig, 1, 0) %>% coalesce(0)) %>%
  mutate(flag_cum = factor(cumsum(flag))) %>%
  filter(sig == TRUE) %>%
  group_by(flag_cum) %>%
  summarize(start = min(hours), end = max(hours)) %>%
  select(-flag_cum)
```

##### Generate model predictions

For visualization, we will generate predictions from each model.

```r
# All care
gam_00.pred <- predict(gam_00, newdata = nd, se.fit = TRUE) %>%
  as_tibble() %>%
  bind_cols(nd) %>%
  select(hours, trt, fit, se.fit) %>%
```

```r
  # bounds = +/- 2 standard errors
  mutate(lower = fit - (2*se.fit),
         upper = fit + (2*se.fit),
         # back-transform to response scale
         fit.response = inv_logit(fit),
         lower.response = inv_logit(lower),
         upper.response = inv_logit(upper))

# MPG care
gam_10.pred <- predict(gam_10, newdata = nd, se.fit = TRUE) %>%
  as_tibble() %>%
  bind_cols(nd) %>%
  select(hours, trt, fit, se.fit) %>%
  # bounds = +/- 2 standard errors
  mutate(lower = fit - (2*se.fit),
         upper = fit + (2*se.fit),
         # back-transform to response scale
         fit.response = inv_logit(fit),
         lower.response = inv_logit(lower),
         upper.response = inv_logit(upper))
```

#### Visualize

These plots show the predicted probability of receiving care through time and across treatments, accounting for the fixed intercept effect of treatment but omitting the random effects of colony and site. The predicted mean response is depicted as a line with an uncertainty band corresponding to 2 x the standard error of the estimate. Intervals during which the probability of receiving care differs across treatments ($p < 0.05$) are highlighted with gray bands.

```r
# Total care
ggplot(gam_00.pred, aes(hours, fit.response, color = trt, fill = trt)) +
  # shade regions where smooth contrast is significant
  geom_rect(data = gam_00_contrast_intervals,
            aes(xmin = start,
                xmax = end,
                ymin = -Inf,
                ymax = Inf),
            inherit.aes = FALSE,
            fill = "gray80") +
  geom_ribbon(aes(ymin = lower.response, ymax = upper.response),
              alpha = 0.4, color = NA) +
  geom_line() +
  labs(x = "Time [h]", y = "Probability of receiving wound care",
       color = "Treatment", fill = "Treatment") +
  scale_x_continuous(breaks = c(0,2,4,6,8,10,12,14,16,18,20,22,24)) +
  scale_y_continuous(breaks = c(0,0.25,0.5,0.75,1), limits = c(0, 1)) +
  theme_classic() +
  theme(legend.position = c(0.87, 0.85))
```

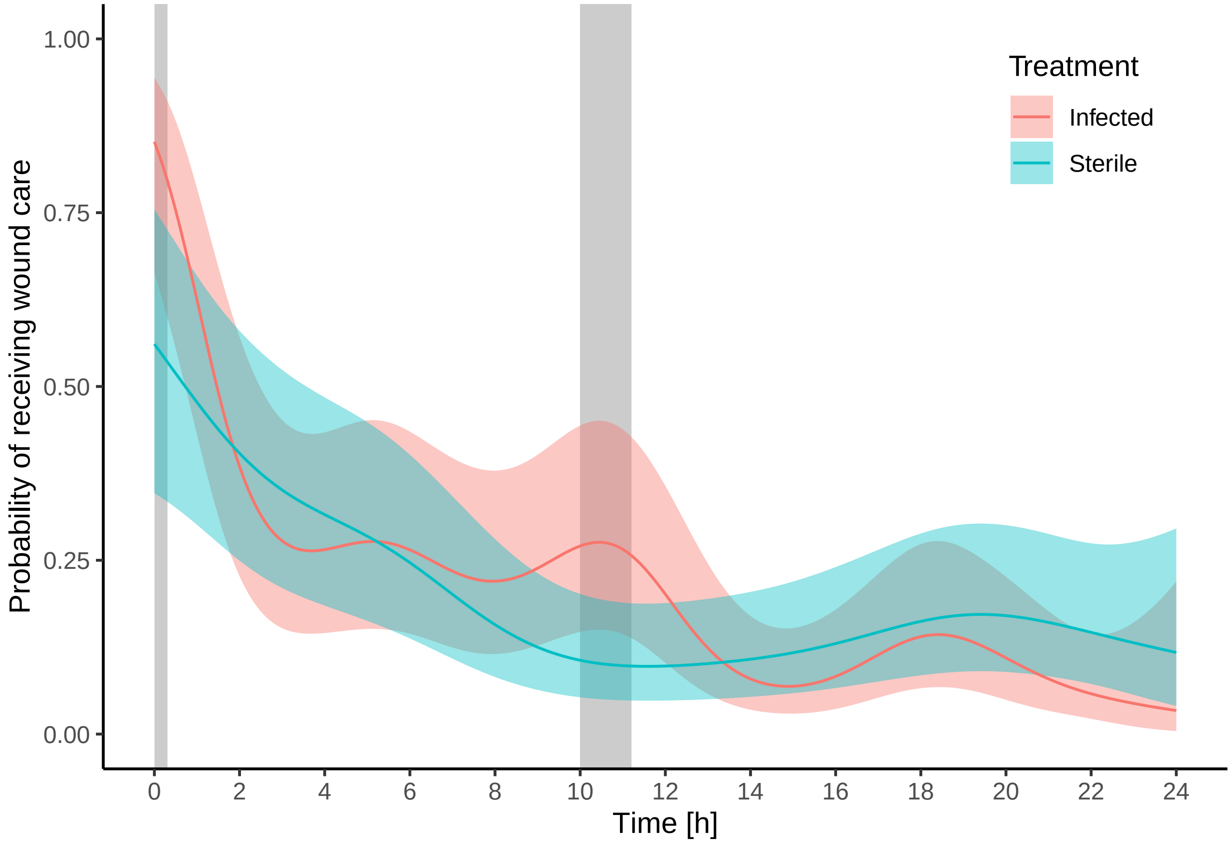

```
# MPG care
ggplot(gam_10.pred, aes(hours, fit.response, color = trt, fill = trt)) +
  # shade regions where smooth contrast is significant
  geom_rect(data = gam_10_contrast_intervals,
            aes(xmin = start,
                xmax = end,
                ymin = -Inf,
                ymax = Inf),
            inherit.aes = FALSE,
            fill = "gray80") +
  geom_ribbon(aes(ymin = lower.response, ymax = upper.response),
              alpha = 0.4, color = NA) +
  geom_line() +
  labs(x = "Time [h]", y = "Probability of receiving MG wound care",
       color = "Treatment", fill = "Treatment") +
  scale_x_continuous(breaks = c(0,2,4,6,8,10,12,14,16,18,20,22,24)) +
  scale_y_continuous(breaks = c(0,0.25,0.5,0.75,1), limits = c(0, 1)) +
  theme_classic() +
  theme(legend.position = c(0.87, 0.85))
```

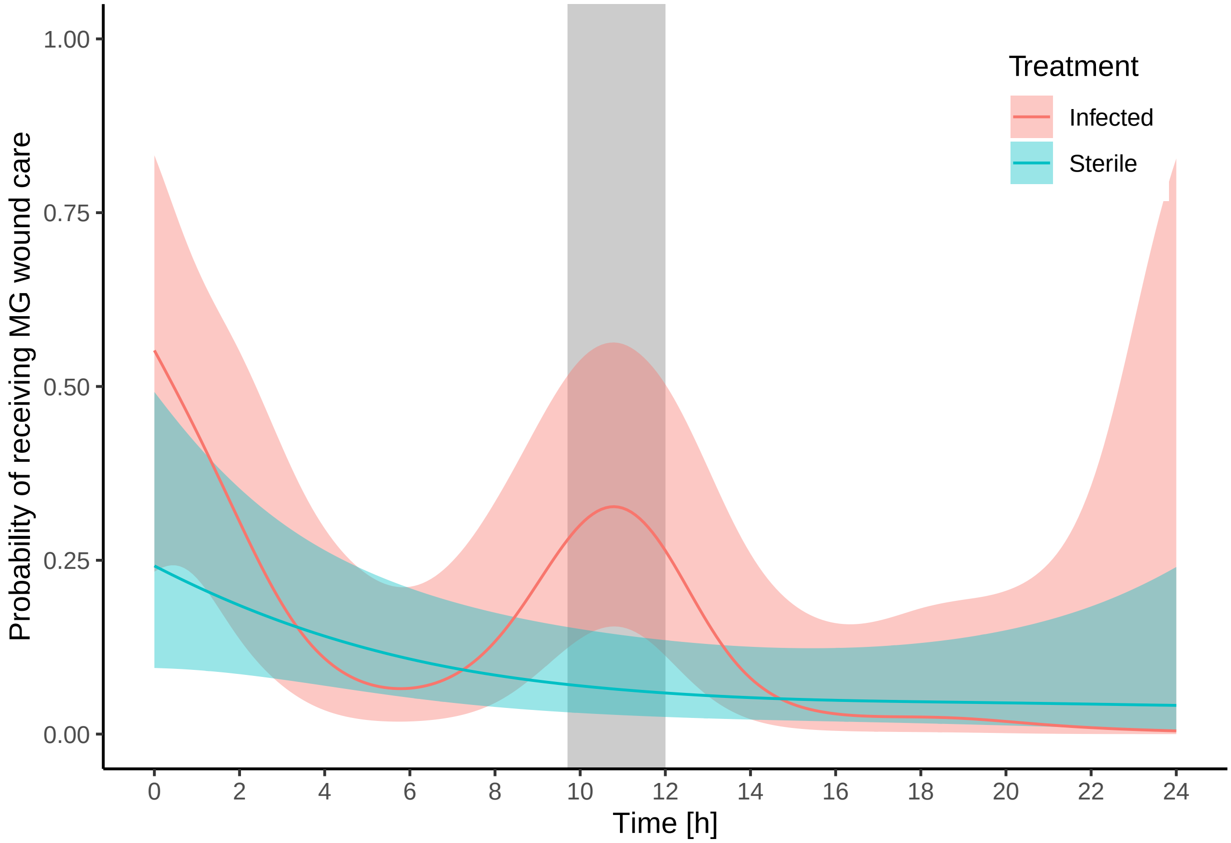
