## Supplemental Tables 1-13 for "Antimicrobial wound care in an ant society"

### Antimicrobial woundcare in an ant society

**This PDF file includes:**

Supplemental Data Tables 1 to 13
Captions for Supplemental Movies 1 to 2

**Other Supplementary Materials for this manuscript include the following:**

Supplemental Movies 1 to 2
Supplemental Model Workflow

| State 1 | State 2 | *t ratio* | *P* |
|---|---|---|---|
| 2h – Sterile | 11h – Sterile | -1.54 | 0.26 |
| 2h – Sterile | 2h – Infected | -3.08 | 0.012* |
| 2h – Sterile | 11h – Infected | -6.75 | <0.001*** |
| 11h – Sterile | 2h – Infected | 1.54 | 0.26 |
| 11h – Sterile | 11h – Infected | -5.21 | <0.001*** |
| 2h – Infected | 11h – Infected | -3.66 | 0.003** |

**Supplemental Table 1 | Statistical differences in bacterial load (16S gene copies) between the treatments presented in Fig 1a.** Linear mixed effect model (Random Factor: Colony: Variance=0, Std. Dev.=0; Residual: Variance=2.26, Std. Dev.=1.50; Likelihood ratio test of model vs intercept only model: $X^2_3$= 34.9, $P$<0.001), $n$=10 per group. Post-hoc analysis with least square means with Holm-Bonferroni correction for the 4 groups.

| **Permutation t-test results at 2 hours** | | | |
|---|---|---|---|
| **Pathogen** | **$R^2$** | ***t*** | ***P*** |
| *Acinetobacter* | 0.15 | -1.76 | 0.09 |
| *Bacillus* | 0.09 | -1.32 | 0.2 |
| *Burkholderia* | 0.97 | 23.89 | <0.001*** |
| *Corynebacterium* | 0.43 | 3.68 | 0.002** |
| *Floricoccus* | 0.01 | -0.48 | 0.6 |
| *Gordonia* | 0.01 | -0.37 | 0.7 |
| *Klebsiella* | 0.92 | 14.73 | <0.001*** |
| *Corynebacteriales* | 0.02 | 0.62 | 0.5 |
| *Pseudomonas* | 0.35 | 3.11 | 0.005** |
| *Streptomyces* | 0.56 | -4.79 | <0.001*** |
| *Tsukamurella* | 0.03 | -0.78 | 0.4 |

| **Permutation t-test results at 11 hours** | | | |
|---|---|---|---|
| **Pathogen** | **$R^2$** | ***t*** | ***P*** |
| *Acinetobacter* | 0.07 | -1.18 | 0.26 |
| *Bacillus* | 0.08 | -1.26 | 0.2 |
| *Burkholderia* | 0.98 | 27.61 | <0.001*** |
| *Corynebacterium* | 0.02 | 0.60 | 0.56 |
| *Floricoccus* | 0.01 | -0.40 | 0.7 |
| *Gordonia* | 0.27 | -2.58 | 0.02* |
| *Klebsiella* | 0.81 | 8.65 | <0.001*** |
| *Corynebacteriales* | <0.01 | -0.03 | 0.97 |
| *Pseudomonas* | 0.28 | 2.62 | 0.02* |
| *Streptomyces* | 0.19 | -2.04 | 0.08 |
| *Tsukamurella* | 0.01 | 0.31 | 0.8 |

**Supplemental Table 2 | Statistical differences between infected and sterile ants in pathogen abundance for the different bacterial genera at 2 and 11 hours presented in Fig. 1b.** Statistical results calculated with a permutation t-test with Holm-Bonferroni correction.

**Mixed effects Cox Model**

| Experiment | Exp(coef) | *Z* | *P* |
|---|---|---|---|
| Nest – Sterile | 0.47 | -1.04 | 0.3 |
| Nest – Infected | 0.83 | -1.32 | 0.75 |
| Isolation – Infected | 4.58 | 4.25 | <0.001*** |

**Least square means (post-hoc analysis)**

| Experiment 1 | Experiment 2 | *Z* | *P* |
|---|---|---|---|
| Isolation – Sterile | Nest – Sterile | 1.040 | 0.89 |
| Isolation – Sterile | Nest – Infected | 0.324 | 1 |
| Isolation – Sterile | Isolation – Infected | -4.246 | <0.001*** |
| Nest – Sterile | Nest – Infected | -0.630 | 1 |
| Nest – Sterile | Isolation – Infected | -2.916 | 0.017* |
| Nest – Infected | Isolation – Infected | -2.759 | 0.023* |

**Supplemental Table 3 | Statistical differences for mortality between the different treatments presented in Fig 1c.** Mixed effects Cox proportional hazards regression model: Random factor: Colony: Variance: 0.11, Std. Dev. 0.33. Likelihood ratio test of model vs intercept only model: $X^2_3$= 16.4, $P$<0.001. Post-hoc analysis with least square means with Holm-Bonferroni correction for the 4 groups.

| Mixed effects Cox Model | | | |
|---|---|---|---|
| **Experiment** | **Exp(coef)** | ***Z*** | ***P*** |
| Nest – Sterile | 0.43 | -1.14 | 0.25 |
| Nest – Infected | 0.20 | -1.59 | 0.11 |
| Isolation – Infected | 4.22 | 4.20 | <0.001*** |

| Least square means (post-hoc analysis) | | | |
|---|---|---|---|
| **Experiment 1** | **Experiment 2** | ***Z*** | ***P*** |
| Isolation – Sterile | Nest – Sterile | 1.145 | 0.50 |
| Isolation – Sterile | Nest – Infected | 1.590 | 0.34 |
| Isolation – Sterile | Isolation – Infected | -4.197 | <0.001*** |
| Nest – Sterile | Nest – Infected | 0.638 | 0.52 |
| Nest – Sterile | Isolation – Infected | -2.994 | 0.014* |
| Nest – Infected | Isolation – Infected | -2.943 | 0.014* |

**Supplemental Table 4 | Statistical differences for mortality between the different treatments presented in Fig 2a.** Mixed effects Cox proportional hazards regression model: Random factor: Colony: Variance: 0.45, Std. Dev. 0.66. Likelihood ratio test of model vs intercept only model: $X^2_3$= 24.59, $P$<0.001. Post-hoc analysis with least square means with Holm-Bonferroni correction for the 4 groups.

| State 1 | State 2 | *t* ratio | *P* |
|---|---|---|---|
| Nest – Sterile – 2h | Nest – Infected – 2h | 0.061 | 1 |
| Nest – Sterile – 2h | Isolation – Sterile – 2h | 0.043 | 1 |
| Nest – Sterile – 2h | Isolation – Infected – 2h | 0.056 | 1 |
| Nest – Sterile – 2h | Nest – Sterile – 11h | 0.039 | 1 |
| Nest – Sterile – 2h | Nest – Infected – 11h | 0.024 | 1 |
| Nest – Sterile – 2h | Isolation – Sterile – 11h | 0.017 | 1 |
| Nest – Sterile – 2h | Isolation – Infected – 11h | -4.776 | <0.001*** |
| Nest – Infected – 2h | Isolation – Sterile – 2h | -0.018 | 1 |
| Nest – Infected – 2h | Isolation – Infected – 2h | -0.005 | 1 |
| Nest – Infected – 2h | Nest – Sterile – 11h | -0.022 | 1 |
| Nest – Infected – 2h | Nest – Infected – 11h | -0.037 | 1 |
| Nest – Infected – 2h | Isolation – Sterile – 11h | -0.044 | 1 |
| Nest – Infected – 2h | Isolation – Infected – 11h | -4.837 | <0.001*** |
| Isolation – Sterile – 2h | Isolation – Infected – 2h | 0.013 | 1 |
| Isolation – Sterile – 2h | Nest – Sterile – 11h | -0.004 | 1 |
| Isolation – Sterile – 2h | Nest – Infected – 11h | -0.019 | 1 |
| Isolation – Sterile – 2h | Isolation – Sterile – 11h | -0.026 | 1 |
| Isolation – Sterile – 2h | Isolation – Infected – 11h | -4.819 | <0.001*** |
| Isolation – Infected – 2h | Nest – Sterile – 11h | -0.017 | 1 |
| Isolation – Infected – 2h | Nest – Infected – 11h | -0.032 | 1 |
| Isolation – Infected – 2h | Isolation – Sterile – 11h | -0.039 | 1 |
| Isolation – Infected – 2h | Isolation – Infected – 11h | -4.832 | <0.001*** |
| Nest – Sterile – 11h | Nest – Infected – 11h | -0.015 | 1 |
| Nest – Sterile – 11h | Isolation – Sterile – 11h | -0.023 | 1 |
| Nest – Sterile – 11h | Isolation – Infected – 11h | -4.816 | <0.001*** |
| Nest – Infected – 11h | Isolation – Sterile – 11h | -0.007 | 1 |
| Nest – Infected – 11h | Isolation – Infected – 11h | -4.800 | <0.001*** |
| Isolation – Sterile – 11h | Isolation – Infected – 11h | -4.793 | <0.001*** |

**Supplemental Table 5 | Statistical differences in bacterial load (*P. aeruginosa*) between the treatments presented in Fig 2b.** Linear mixed effect model (Random Factor: Colony: Variance=$3.5*10^{-13}$, Std. Dev.=$5.8*10^{-7}$; Residual: Variance=50590, Std. Dev.=224.9; Likelihood ratio test of model vs intercept only model: $X^2_9$= 36.3, *P*<0.001), *n*=6 per group. Post-hoc analysis with least square means with Holm-Bonferroni correction for the 8 groups.

| Mixed effects Cox Model | | | |
|---|---|---|---|
| **Experiment** | **Exp(coef)** | ***Z*** | ***P*** |
| Isolation – clotted – Infected | 9.07 | 3.13 | <0.001*** |
| Isolation – open – Sterile | 1.32 | 0.46 | 0.65 |
| Isolation – open – Infected | 9.70 | 3.82 | <0.001*** |
| Nest – clotted – Sterile | 0.30 | -1.05 | 0.29 |
| Nest – clotted – Infected | 7.09 | 3.02 | 0.003** |
| Nest – open – Sterile | 0.29 | -1.07 | 0.28 |
| Nest – open – Infected | 1.02 | 0.02 | 0.98 |

| Least square means (post-hoc analysis) | | | |
|---|---|---|---|
| **Experiment 1** | **Experiment 2** | ***Z*** | ***P*** |
| Isolation – clotted – Sterile | Isolation – clotted – Infected | -3.387 | 0.015* |
| Isolation – clotted – Sterile | Isolation – open – Sterile | -0.455 | 1 |
| Isolation – clotted – Sterile | Isolation – open – Infected | -3.818 | 0.003** |
| Isolation – clotted – Sterile | Nest – clotted – Sterile | 1.050 | 1 |
| Isolation – clotted – Sterile | Nest – clotted – Infected | -3.016 | 0.039* |
| Isolation – clotted – Sterile | Nest – open – Sterile | 1.075 | 1 |
| Isolation – clotted – Sterile | Nest – open – Infected | -0.020 | 1 |
| Isolation – clotted – Infected | Isolation – open – Sterile | 5.540 | <0.001*** |
| Isolation – clotted – Infected | Isolation – open – Infected | -0.210 | 1 |
| Isolation – clotted – Infected | Nest – clotted – Sterile | 3.271 | 0.019* |
| Isolation – clotted – Infected | Nest – clotted – Infected | 0.603 | 1 |
| Isolation – clotted – Infected | Nest – open – Sterile | 3.298 | 0.019* |
| Isolation – clotted – Infected | Nest – open – Infected | 3.364 | 0.015* |
| Isolation – open – Sterile | Isolation – open – Infected | -8.780 | <0.001*** |
| Isolation – open – Sterile | Nest – clotted – Sterile | 1.464 | 1 |
| Isolation – open – Sterile | Nest – clotted – Infected | -4.870 | <0.001*** |
| Isolation – open – Sterile | Nest – open – Sterile | 1.493 | 1 |
| Isolation – open – Sterile | Nest – open – Infected | 0.429 | 1 |
| Isolation – open – Infected | Nest – clotted – Sterile | 3.447 | 0.012* |
| Isolation – open – Infected | Nest – clotted – Infected | 0.983 | 1 |
| Isolation – open – Infected | Nest – open – Sterile | 3.476 | 0.012* |
| Isolation – open – Infected | Nest – open – Infected | 3.793 | 0.004** |
| Nest – clotted – Sterile | Nest – clotted – Infected | -3.038 | 0.038* |
| Nest – clotted – Sterile | Nest – open – Sterile | 0.021 | 1 |
| Nest – clotted – Sterile | Nest – open – Infected | -1.064 | 1 |
| Nest – clotted – Infected | Nest – open – Sterile | 3.065 | 0.037* |
| Nest – clotted – Infected | Nest – open – Infected | 2.992 | 0.039* |
| Nest – open – Sterile | Nest – open – Infected | -1.089 | 1 |

**Supplemental Table 6 | Statistical differences for mortality between the different treatments presented in Extended Data Fig. 5.** Mixed effects Cox proportional hazards regression model: Random factor: Colony: Variance: 0.00008, Std. Dev. 0.0089. Likelihood ratio test of model vs intercept only model: $X^2_7$= 119.9, $P$<0.001. Post-hoc analysis with least square means with Holm-Bonferroni correction for the 8 groups (28 tests). Nest: focal ant kept in sub-colony; Isolation: focal ant kept alone; clotted: MG clotted with acrylic color both in the focal ant and nestmates; open: MG was kept open both in the focal ant and nestmates; Sterile: wound of focal ant was exposed to PBS; Infected: wound of focal ant was exposed to 0.05 OD of *P. aeruginosa*.

| **Group 1** | **Group 2** | $R^2$ | *F* | *P* |
|---|---|---|---|---|
| Infected – 0h | Infected – 2h | 0.13 | 2.42 | 0.05* |
| Infected – 0h | Infected – 11h | 0.19 | 3.82 | 0.008** |
| Infected – 0h | Sterile – 0h | 0.13 | 0.13 | 0.96 |
| Infected – 0h | Sterile – 2h | 0.11 | 1.95 | 0.09 |
| Infected – 0h | Sterile – 11h | 0.06 | 0.96 | 0.34 |
| Infected – 2h | Infected – 11h | 0.063 | 1.48 | 0.13 |
| Infected – 2h | Sterile – 0h | 0.17 | 3.26 | 0.005** |
| Infected – 2h | Sterile – 2h | 0.008 | 0.17 | 0.87 |
| Infected – 2h | Sterile – 11h | 0.13 | 3.40 | 0.03* |
| Infected – 11h | Sterile – 0h | 0.23 | 4.90 | 0.001*** |
| Infected – 11h | Sterile – 2h | 0.07 | 1.56 | 0.16 |
| Infected – 11h | Sterile – 11h | 0.19 | 5.4 | 0.007** |
| Sterile – 0h | Sterile – 2h | 0.15 | 2.74 | 0.03* |
| Sterile – 0h | Sterile – 11h | 0.03 | 0.53 | 0.45 |
| Sterile – 2h | Sterile – 11h | 0.14 | 3.54 | 0.04* |

**Supplemental Table 7 | Statistical differences in CHC-profile composition between the treatments.** Permutational multivariate analysis of variance using Bray-Curtis dissimilarity matrices (ADONIS: formula: CHCdistancematrix ~ Treatment&Time*Sociality with Colony as random factor). Treatment&Time: Df=5, $R^2$=0.18, *F*=2.32, *P*=0.009; Sociality: Df=1, $R^2$=0.012, *F*=0.77, *P*=0.47; Treatment&Time*Sociality: Df=3, $R^2$=0.04, *F*=0.86, *P*=0.54; *n*=12 per group, except for timepoint 0: *n*=6. Post-hoc analysis with a pairwise ADONIS with colony as random factor and correction for multiple comparisons across the 6 groups.

| Compound | Compound type | Ret. Index | Sterile 0h *n*=6 | Sterile 2h *n*=12 | Sterile 11h *n*=12 | Infected 0h *n*=6 | Infected 2h *n*=12 | Infected 11h *n*=12 |
|---|---|---|---|---|---|---|---|---|
| C20 | Alkane | 2000 | 0.0±0.1 | 0.1±0.1 | 0.1±0.1 | 0.0±0.0 | 0.0±0.0 | 0.1±0.0 |
| C21 | Alkane | 2100 | 1.0±0.5 | 1.0±0.4 | 1.6±0.8 | 1.1±0.8 | 0.9±0.4 | 0.9±0.3 |
| 10-MeC21 | Methyl | 2140 | 0.6±0.1 | 0.5±0.2 | 0.5±0.2 | 0.4±0.1 | 0.4±0.2 | 0.5±0.2 |
| 3-MeC21 | Methyl | 2173 | 0.1±0.1 | 0.1±0.1 | 0.1±0.1 | 0.0±0.0 | 0.0±0.1 | 0.1±0.1 |
| C22 | Alkane | 2200 | 0.4±0.4 | 0.3±0.1 | 0.4±0.1 | 0.4±0.6 | 0.3±0.1 | 0.4±0.2 |
| C23en_1 | Alkene | 2273 | 1.6±1.4 | 1.7±0.9 | 1.3±0.7 | 1.7±1.8 | 1.4±0.5 | 1.5±0.8 |
| C23en_2 | Alkene | 2280 | 0.0±0.0 | 0.0±0.0 | 0.0±0.0 | 0.0±0.0 | 0.0±0.0 | 0.0±0.1 |
| C23 | Alkane | 2300 | 9.2±3.0 | 6.8±1.3 | 7.3±1.4 | 9.5±2.8 | 6.8±1.4 | 6.5±1.1 |
| 11-MeC23 | Methyl | 2335 | 1.2±0.6 | 1.3±0.2 | 1.2±0.4 | 1.2±0.7 | 1.1±0.3 | 1.2±0.3 |
| 5-MeC23 | Methyl | 2353 | 0.2±0.2 | 0.2±0.1 | 0.2±0.1 | 0.1±0.1 | 0.2±0.1 | 0.2±0.1 |
| 3-MeC23 | Methyl | 2372 | 0.5±0.5 | 0.2±0.3 | 0.2±0.3 | 0.1±0.1 | 0.0±0.0 | 0.4±0.1 |
| C24 | Alkane | 2400 | 1.3±0.1 | 1.1±0.2 | 1.4±0.2 | 1.1±0.1 | 1.2±0.2 | 1.2±0.2 |
| 3,7-DiMeC23 | Dimethyl | 2409 | 0.0±0.0 | 0.0±0.0 | 0.1±0.2 | 0.0±0.0 | 0.0±0.1 | 0.2±0.1 |
| C25en_1 | Alkene | 2471 | 0.5±0.4 | 0.6±0.4 | 0.5±0.3 | 0.5±0.7 | 0.5±0.2 | 0.7±0.4 |
| C25en_2 | Alkene | 2478 | 0.0±0.0 | 0.1±0.1 | 0.1±0.1 | 0.0±0.0 | 0.0±0.1 | 0.1±0.1 |
| C25 | Alkane | 2500 | 29.8±7.9 | 22.0±4.5 | 24.4±11.0 | 23.6±12.1 | 22.6±3.6 | 19.7±5.0 |
| 11-;13-MeC25 | Methyl | 2531 | 0.2±0.1 | 0.2±0.1 | 0.3±0.1 | 0.2±0.1 | 0.2±0.0 | 0.3±0.0 |
| 3-MeC25 | Methyl | 2572 | 0.3±0.2 | 0.2±0.0 | 0.2±0.1 | 0.1±0.1 | 0.2±0.1 | 0.2±0.1 |
| C26 | Alkane | 2600 | 0.4±0.2 | 0.7±0.2 | 0.7±0.2 | 0.5±0.1 | 0.7±0.1 | 0.7±0.2 |
| 13-;11-;9-MeC26 | Methyl | 2614 | 0.0±0.0 | 0.0±0.0 | 0.0±0.0 | 0.0±0.0 | 0.0±0.0 | 0.1±0.1 |
| C27en | Alkene | 2676 | 1.2±0.4 | 1.7±0.4 | 1.4±0.3 | 1.4±1.0 | 1.7±0.4 | 1.8±0.3 |
| C27 | Alkane | 2700 | 4.3±1.5 | 3.5±0.9 | 4.8±0.7 | 3.7±1.9 | 3.9±1.2 | 3.9±1.0 |
| 13-;11-MeC27 | Methyl | 2732 | 0.1±0.1 | 0.1±0.2 | 0.2±0.2 | 0.0±0.0 | 0.1±0.2 | 0.1±0.2 |
| C29en_1 | Alkene | 2870 | 0.0±0.0 | 0.0±0.0 | 0.0±0.0 | 0.0±0.0 | 0.0±0.0 | 0.1±0.1 |
| C29en_2 | Alkene | 2878 | 1.3±0.7 | 1.5±0.4 | 1.5±0.8 | 1.4±0.4 | 1.5±0.4 | 1.7±0.7 |
| C29 | Alkane | 2900 | 0.8±0.2 | 1.0±0.3 | 1.1±0.2 | 0.9±0.2 | 1.0±0.2 | 1.2±0.3 |
| 15-;13-;11-MeC29 | Methyl | 2933 | 0.0±0.0 | 0.0±0.0 | 0.0±0.1 | 0.0±0.0 | 0.0±0.0 | 0.2±0.2 |
| C30en | Alkene | 2980 | 0.0±0.0 | 0.2±0.2 | 0.1±0.1 | 0.0±0.0 | 0.2±0.1 | 0.2±0.1 |
| C31dien | Alkadiene | 3050 | 0.6±0.2 | 1.0±0.3 | 0.7±0.5 | 0.5±0.3 | 0.8±0.3 | 1.2±0.5 |
| C31en_1 | Alkene | 3070 | 0.4±0.5 | 0.7±0.2 | 0.6±0.4 | 0.3±0.4 | 0.8±0.3 | 0.9±0.3 |
| C31en_2 | Alkene | 3084 | 19.4±3.8 | 20.8±1.0 | 18.5±2.2 | 19.9±4.9 | 21.0±0.9 | 19.8±1.1 |
| C31 | Alkane | 3100 | 0.0±0.0 | 0.0±0.0 | 0.3±0.4 | 0.2±0.3 | 0.5±0.2 | 0.6±0.0 |
| 15-;13-;11-MeC31 | Methyl | 3130 | 0.0±0.0 | 0.1±0.2 | 0.1±0.1 | 0.0±0.0 | 0.1±0.1 | 0.2±0.2 |
| C32dien | Alkadiene | 3148 | 0.1±0.2 | 0.3±0.3 | 0.3±0.2 | 0.0±0.0 | 0.4±0.2 | 0.5±0.3 |
| C33trien | Alkatriene | 3232 | 0.0±0.0 | 0.1±0.0 | 0.0±0.0 | 0.0±0.0 | 0.1±0.1 | 0.1±0.0 |
| C33dien_1 | Alkadiene | 3247 | 23.9±2.4 | 28.8±1.8 | 24.6±2.7 | 23.9±6.4 | 27.9±2.5 | 27.6±2.5 |
| C33dien_2 | Alkadiene | 3274 | 0.0±0.0 | 0.0±0.0 | 0.0±0.0 | 0.0±0.0 | 0.1±0.1 | 0.4±0.5 |
| C35trien | Alkatriene | 3449 | 0.0±0.0 | 0.0±0.0 | 0.0±0.0 | 0.0±0.0 | 0.1±0.1 | 0.2±0.1 |
| C35dien | Alkadiene | 3472 | 0.0±0.0 | 0.0±0.0 | 0.0±0.0 | 0.0±0.0 | 0.0±0.0 | 0.1±0.1 |

**Supplemental Table 8 | Chemical composition of the cuticular hydrocarbon profiles.** Median relative percentages with median absolute deviation of all cuticular hydrocarbons between infected and sterile ants across the three timepoints (0h, 2h, 11h).

| Group 1 | Group 2 | *P* adj. |
|---|---|---|
| Sterile – 0h | Sterile – 2h | 0.01* |
| Sterile – 0h | Sterile – 11h | 0.92 |
| Sterile – 0h | Infected – 0h | 0.99 |
| Sterile – 0h | Infected – 2h | 0.02* |
| Sterile – 0h | Infected – 11h | 0.002** |
| Sterile – 2h | Sterile – 11h | 0.04* |
| Sterile – 2h | Infected – 0h | 0.07 |
| Sterile – 2h | Infected – 2h | 1 |
| Sterile – 2h | Infected – 11h | 0.98 |
| Sterile – 11h | Infected – 0h | 1 |
| Sterile – 11h | Infected – 2h | 0.049* |
| Sterile – 11h | Infected – 11h | 0.005** |
| Infected – 0h | Infected – 2h | 0.08 |
| Infected – 0h | Infected – 11h | 0.01* |
| Infected – 2h | Infected – 11h | 0.96 |

**Supplemental Table 9 | Statistical differences for alkadienes between the treatments presented in Extended Data Fig. 6.** AOV model (alkanes: Df=5, Sum Sq=713.6, $F$=2.99, $P$=0.02 Residual: Df=53, Sum Sq=2522; alkenes: Df=5, Sum Sq=124.6, $F$=2.46, $P$=0.04; Residual: Df=53, Sum Sq=536; alkadienes: Df=5, Sum Sq=384.7, $F$=6.82, $P$<0.001 Residual: Df=53, Sum Sq=598; methyl-branched-alkanes: Df=5, Sum Sq=4.13, $F$=0.56, $P$=0.73 Residual: Df=53, Sum Sq=78) $n$=12 per group, except for timepoint zero were $n$=6 for sterile and infected ants. Post-hoc analysis with Tukey Honest Significant differences test. Even though the AOV was significant for alkanes and alkenes, the posthoc analysis with Holm-Bonferroni corrections for multiple testing across the 6 groups did not result in any pairwise significances, the detailed posthoc results are thus only given for the alkadienes.

| Gene IDs | Log2FoldChange | *P* adj. | Time | Putative Function | Biological Process (GO) |
|---|---|---|---|---|---|
| gene_22863 | 1.00068922 | <0.001 | 2h | phosphatidylinositol 4-kinase beta | lipid biosynthetic process |
| gene_30894 | 2.36850382 | <0.001 | 2h | glucose dehydrogenase | glucose metabolic process |
| gene_02329 | -1.037557432 | 0.003 | 11h | endocuticle structural glycoprotein SgAbd-4 | structural constituent of cuticle |
| gene_38881 | -3.401621515 | 0.029 | 11h | sodium-coupled monocarboxylate transporter 1-like | short-chain fatty acid import |
| gene_08375 | -1.532425715 | <0.001 | 11h | lipid storage droplets surface-binding protein 2-like | regulation of lipid storage |
| gene_08376 | 2.55138413 | 0.048 | 11h | lipid storage droplets surface-binding protein 1 | regulation of lipid storage |
| gene_31650 | -2.305854998 | <0.001 | 11h | palmitoyltransferase | acyltransferase activity |
| gene_34515 | -1.801145986 | 0.003 | 11h | very long-chain-fatty-acid--CoA ligase bubblegum | long-chain fatty acid-CoA elongase activity |
| gene_40960 | 3.473641627 | 0.018 | 11h | elongation of very long chain fatty acids protein 1-like | fatty acid elongation |
| gene_14544 | -4.357452189 | 0.002 | 11h | Cuticle protein 6 | structural constituent of cuticle |
| gene_02681 | 1.051188424 | <0.001 | 11h | phospholipase A2-like | lipid metabolic process |
| gene_14361 | -1.56417983 | 0.045 | 11h | pancreatic triacylglycerol lipase | lipid metabolic process |
| gene_23311 | 1.428011778 | 0.027 | 11h | microsomal triglyceride transfer protein | lipid metabolic process |
| gene_29857 | 1.734871622 | 0.030 | 11h | diacylglycerol kinase eta | lipid metabolic process |
| gene_39381 | -2.135617551 | 0.039 | 11h | alkaline ceramidase | lipid metabolic process |
| gene_18230 | -0.613998454 | 0.004 | 11h | sterol O-acyltransferase | fatty-acyl-CoA binding |
| gene_39586 | -0.944340779 | 0.012 | 11h | UDP-xylose and UDP-N-acetylglucosamine transporter-like | carbohydrate transport |
| gene_38486 | 2.738654673 | <0.001 | 11h | sorbitol dehydrogenase-like | carbohydrate metabolism |
| gene_18615 | -1.748361395 | <0.001 | 11h | hydroxymethylglutaryl-CoA synthase 1 | acetyl-CoA metabolic process |
| gene_38395 | -1.037557432 | 0.045 | 11h | acid phosphatase | phosphoric ester hydrolase activity |

**Supplemental Table 10 | List of genes putatively implicated in CHC production and lipid metabolism being differentially expressed between sterile and infected ants presented in Extended Data Fig. 7.** Positive Log2FoldChange values correspond to genes up-regulated in infected ants. *P*-values are corrected for multiple testing using the Benjamini and Hochberg method.

| Gene IDs | Log2FoldChange | *P* adj. | Time | Putative Function | Biological Process (GO) |
|---|---|---|---|---|---|
| gene_02403 | 2.35332709 | 0.001 | 2h | tyrosine kinase receptor Cad96Ca-like | positive regulation of wound healing |
| gene_07900 | 1.07975465 | 0.009 | 2h | peptidoglycan-recognition protein SC2-like PGRP-LB | positive regulation of Toll signaling pathway |
| gene_12381 | 5.27341349 | <0.001 | 2h | Pirk | negative regulation of peptidoglycan recognition protein signaling pathway |
| gene_14574 | 2.45236598 | 0.001 | 2h | Abaecin | hemolymph coagulation |
| gene_25734 | 5.05363939 | <0.001 | 2h | Hymenoptaecin | innate immune response |
| gene_25736 | 4.00166659 | <0.001 | 2h | Hymenoptaecin | innate immune response |
| gene_39135 | 1.3685947 | 0.003 | 2h | serine protease inhibitor 88Ea-like | negative regulation of innate immune response (Toll) |
| gene_01714 | 2.68170259 | 0.006 | 11h | PDGF- and VEGF-related factor 3 | hemocyte migration |
| gene_07900 | 2.55345461 | <0.001 | 11h | peptidoglycan-recognition protein SC2-like PGRP-LB | positive regulation of Toll signaling pathway |
| gene_11103 | 0.85188693 | 0.006 | 11h | fas-associated death domain protein (Fadd) | positive regulation of innate immune response |
| gene_12381 | 5.63944847 | <0.001 | 11h | Pirk | negative regulation of peptidoglycan recognition protein signaling pathway |
| gene_14574 | 4.429955 | <0.001 | 11h | Abaecin | hemolymph coagulation |
| gene_17612 | 1.56736442 | <0.001 | 11h | phenoloxidase-activating factor 2 (PPAF2) | innate immune response |
| gene_25734 | 7.21006972 | <0.001 | 11h | Hymenoptaecin | innate immune response |
| gene_25736 | 6.31954344 | <0.001 | 11h | Hymenoptaecin | innate immune response |
| gene_25792 | 0.65380865 | 0.021 | 11h | Autophagy-related protein 16-1 (Atg 16) | positive regulation of autophagy |
| gene_29419 | -1.8089618 | 0.003 | 11h | protein croquemort-like | immune response-regulating cell surface receptor signaling pathway involved in phagocytosis |
| gene_30388 | 0.40053527 | 0.009 | 11h | coronin-1C-A (coro) | defence response to fungus |
| gene_34607 | 1.95877062 | <0.001 | 11h | Cytokine receptor | receptor signaling pathway via JAK-STAT |
| gene_37992 | 0.77372933 | <0.001 | 11h | 2-oxoglutarate-dependent dioxygenase htyE (2OGDDs) | biosynthesis of the beta-lactam antibiotics |
| gene_39135 | 2.15522495 | <0.001 | 11h | serine protease inhibitor 88Ea-like | negative regulation of innate immune response (Toll) |
| gene_39979 | 2.55922106 | <0.001 | 11h | serine protease easter (ea) | positive regulation of Toll signaling pathway |
| gene_41380 | 1.49200932 | <0.001 | 11h | serine protease gd | positive regulation of Toll signaling pathway |
| gene_41384 | 2.08260184 | <0.001 | 11h | serine protease gd | positive regulation of Toll signaling pathway |

**Supplemental Table 11 | Immune system related genes differentially expressed between sterile and infected ants presented in Extended Data Fig. 7.** Positive Log2FoldChange values correspond to genes up-regulated in infected ants. *P*-values are corrected for multiple testing using the Benjamini and Hochberg method.

| *D. mel* ortholog | Protein IDs | Function | Orthology Depth | % MG Content | <20 kDa | Implications for wound healing from orthologs |
|---|---|---|---|---|---|---|
| - | gene_35433 | Unknown | None | 13.239 | X | |
| CG15203 | gene_35399 | Unknown | Arthropoda, poss. slime mold | 5.943 | X | Toxin-like, O_Venom |
| Hml | gene_35649 | chymotrypsin inhibitor | Endopterygota | 5.568 | X | O_Hemocyte, O_Venom |
| - | gene_41829 | Unknown | Hymenoptera | 4.237 | X | Toxin-like, O_Venom |
| straw | gene_12257 | laccase | Pterygota | 3.192 | | O_Melanization |
| yellow-d | gene_33077 | MRJP | Hymenoptera | 2.919 | | O_Antimicriobial |
| straw | gene_12258 | laccase | Pterygota | 1.935 | | O_Melanization |
| Gba1b | gene_20433 | glucosylceramidase | Bilateria | 1.077 | | |
| yellow-b | gene_33078 | MRJP | Hymenoptera | 0.797 | | O_Antimicrobial |
| CG34034 | gene_25950 | omega-conotoxin-like | Insecta, poss. bacteria, fungi | 0.751 | X | Toxin-like, O_Venom |
| - | gene_00214 | Unknown | Formicidae | 0.646 | X | Toxin-like, O_Venom |
| - | gene_13637 | Unknown | Formicidae, possibly sawfly | 0.586 | X | |
| CG6426 | gene_37453 | lysozyme | Neoptera | 0.431 | X | O_Antimicrobial |
| - | gene_16798 | kielin/chordin-like | Endopterygota | 0.244 | | |
| - | gene_25492 | Unknown | Formicidae | 0.236 | X | Toxin-like, O_Venom |
| eater | gene_29492 | VWDE / eater / fibrilin | Neoptera | 0.181 | | O_Hemocyte |
| - | gene_25504 | Odorant binding protein, GP9-like | Aculeata | 0.168 | X | |
| - | gene_25951 | omega-conotoxin-like | Insecta, poss. bacteria, fungi | 0.147 | X | Toxin-like, O_Venom |
| - | gene_00213 | Unknown | Formicidae | 0.128 | X | O_Venom |
| crok | gene_28725 | Unknown/quiver | Neoptera | 0.124 | X | O_Venom |
| - | gene_24061 | Unknown | Hymenoptera, poss. bacteria | 0.107 | X | O_Venom |
| CG42259 | gene_35648 | chymotrypsin inhibitor | Invertebrates, fungi, viruses | 0.070 | X | O_Wound_response, Toxin-like, O_Venom |
| CG8369 | gene_09378 | kazal-type proteinase inhibitor / vasotab | Neoptera | 0.051 | X | O_Vasodilator |
| CG9917 | gene_38892 | Interferon-related developmental regulator | Pterygota | 0.043 | | |
| - | gene_20297 | Myrosinase-1-like | Endopterygota | 0.036 | | |
| Nep2 | gene_06446 | Neprilysin / membrane metallo-endopeptidase | Pterygota | 0.034 | | |
| CG30197 | gene_25140 | waprin-Thr1 | Neoptera | 0.027 | X | Toxin-like, O_Venom, O_Antimicrobial |
| CG30197 | gene_25139 | waprin-Phi1 | Pteroygota | 0.023 | | Toxin-like, O_Venom, O_Antimicrobial |
| - | gene_08904 | CREG1-like | Pancrustacea | 0.021 | | |
| CG15140 | gene_38752 | Prisilkin/trithorax/pro-resilin | Endopterygota | 0.021 | | |
| - | gene_16797 | kielin/chordin-like | Endopterygota | 0.020 | | |
| hgo | gene_00180 | homogentisate 1,2-dioxygenase | Bilateria | 0.019 | | O_Melanization |
| CG6414 | gene_06365 | Venom carboxylesterase-6 | Neoptera | 0.014 | | |
| - | gene_00215 | Unknown | Formicidae, possibly bacteria | 0.011 | X | O_Venom |
| l(1)G0289 | gene_07578 | plexin domain-containing protein | Pterygota | 0.011 | | |
| CAH2 | gene_03491 | carbonic anhydrase | Pterygota | 0.008 | | |
| mgl | gene_13013 | sortilin-related receptor | Pterygota | 0.007 | | |
| Alp4 | gene_33376 | alkaline phosphatase | Hexapoda | 0.006 | | |
| Fkbp14 | gene_11011 | FK506 Binding protein, TOR related | Pancrustacea, fungi | 0.006 | X | |
| Atpα | gene_07025 | sodium/potassium-transporting ATPase | Bilateria | 0.004 | | |
| Nrx-1 | gene_10345 | Multiple EGF-like domains / crumbs | Arthropoda | 0.003 | X | Toxin-like |

**Supplemental Table 12 | Proteins from the metapleural gland.** Forty-one proteins had a significantly greater abundance in the metapleural gland atrium than in the hemolymph (see Extended Data Fig. 8b). Abbreviations: X: Low molecular weight (<20kDa); O_: function found in an orthologous protein; *Drosophila melanogaster* orthologs are indicated where there was sufficient similarity. Proteins are sorted by their abundance in the metapleural gland atrium. Function orthology depth (common sequences) and implications were based on protein BLAST hits using the experimental clustered nr database[37].

| Compound | Ret. Index | Relative abundance | Function / Chemical Group |
|---|---|---|---|
| Unidentified_1 | 842 | 0.22±0.26 | NA |
| Furanmethanol | 860 | 0.31±0.15 | Alcohol |
| Similar to 2-Propenamide | 900 | 0.17±0.21 | Amide |
| Similar to 2-Propenamide | 905 | 0.17±0.18 | Amide |
| 2(5H)-Furanone | 918 | 0.31±0.07 | |
| 3-Methyl-2(5H)-furanone | 982 | 0.44±0.18 | |
| Phenol | 993 | 0.21±0.10 | **Acid** |
| Hexanoic acid | 1016 | 0.22±0.12 | **Acid** |
| Unidentified_2 | 1033 | 0.05±0.05 | NA |
| Unidentified_3 | 1049 | 0.13±0.07 | NA |
| 3-Ethyl-2,5-dimethyl-pyrazine | 1083 | 0.57±0.51 | **Alkaloid** |
| Unidentified_4 | 1086 | 0.30±0.09 | NA |
| Unidentified_5 | 1088 | 0.27±0.28 | NA |
| Unidentified_6 | 1091 | 0.39±0.29 | NA |
| Hexanamide | 1129 | 0.06±0.05 | Amide |
| Unidentified_7 | 1143 | 0.40±0.50 | NA |
| 2,3-Dihydro-3,5-dihydoxy-6-methyl-4H-pyran-4-one | 1153 | 0.15±0.15 | |
| Glutarimide | 1160 | 0.13±0.05 | Amide |
| Unidentified_8 | 1174 | 0.09±0.13 | NA |
| Catechol | 1215 | 0.18±0.18 | **Antimicrobial** |
| Unidentified_9 | 1263 | 0.20±0.13 | NA |
| Nonanoic acid | 1288 | 0.42±0.23 | **Acid** |
| Indole | 1299 | 0.93±0.31 | Amine |
| Unidentified_10 | 1303 | 0.39±0.28 | |
| Unidentified_11 | 1310 | 0.30±0.31 | NA |
| 4-Methyl-1,2-benzenediol | 1318 | 0.20±0.26 | NA |
| 2,6-Dimethoxyphenol | 1359 | 0.41±0.38 | |
| Unidentified_12 | 1364 | 0.36±0.23 | NA |
| Unidentified_13 | 1368 | 0.30±0.32 | NA |
| 3-Methyl-indole | 1391 | 0.63±0.17 | **Alkaloid** |
| 5-Oxo-L-proline methyl ester | 1396 | 0.56±0.77 | **Alkaloid** |
| Alkaloid_1 | 1410 | 0.80±0.35 | **Alkaloid** |
| Alkaloid_2 | 1414 | 0.34±0.34 | **Alkaloid** |
| Similar to Gougerotin | 1446 | 0.12±0.12 | **Antibiotic** |
| Caprolactone derivative | 1466 | 0.48±0.39 | |
| Isoxacol derivative | 1470 | 0.17±0.10 | **Antimicrobial component** |
| Unidentified_14 | 1473 | 0.23±0.20 | NA |
| Alkaloid_3 | 1475 | 0.38±0.20 | **Alkaloid** |
| Piperidine derivative_1 | 1479 | 0.63±0.14 | **Alkaloid** |
| Piperidine derivative_2 | 1484 | 0.58±0.33 | **Alkaloid** |
| Piperidine derivative_3 | 1489 | 0.24±0.02 | **Alkaloid** |
| Unidentified_15 | 1511 | 0.12±0.12 | NA |
| Unidentified_16 | 1515 | 0.16±0.23 | NA |
| Pyrimidin derivative | 1520 | 0.11±0.12 | |
| Alkaloid_4 | 1549 | 0.28±0.18 | **Alkaloid** |
| Alkaloid_5 | 1553 | 0.34±0.00 | **Alkaloid** |
| Dodecanoic acid | 1581 | 2.12±1.28 | **Acid** |
| Unidentified_17 | 1587 | 0.40±0.24 | NA |
| Alkaloid_6 | 1617 | 0.42±0.14 | **Alkaloid** |
| Alkaloid_7 | 1626 | 0.68±0.17 | **Alkaloid** |
| Unidentified_18 | 1630 | 0.25±0.31 | NA |
| 12-Hydroxydecanoic acid | 1666 | 0.86±0.79 | **Acid** |
| 10-Hydroxydecanoic acid | 1671 | 2.06±1.91 | **Acid** |
| Alkaloid_8 | 1673 | 0.23±0.22 | **Alkaloid** |
| Deoxyspergualin derivative | 1679 | 0.31±0.30 | **Antibiotic** |

| | | | |
|---|---|---|---|
| Unidentified_19 | 1682 | 0.31±0.12 | NA |
| Unidentified_20 | 1689 | 0.28±0.26 | NA |
| Alkaloid_9 | 1704 | 0.16±0.05 | **Alkaloid** |
| Alkaloid_10 | 1712 | 0.33±0.18 | **Alkaloid** |
| Similar to 3-Methyl-1,4-diazabicyclo[4.3.0]nonan-2,5-dione | 1732 | 3.44±1.12 | |
| Alkaloid_11 | 1739 | 0.69±0.39 | **Alkaloid** |
| Alkaloid_12 | 1747 | 0.29±0.06 | **Alkaloid** |
| Similar to 3-Methyl-1,4-diazabicyclo[4.3.0]nonan-2,5-dione | 1758 | 2.02±0.46 | |
| Alkaloid_13 | 1770 | 0.54±0.38 | **Alkaloid** |
| Alkaloid_14 | 1780 | 3.42±1.30 | **Alkaloid** |
| Alkaloid_15 | 1783 | 1.60±0.72 | **Alkaloid** |
| Alkaloid_16 | 1785 | 0.95±0.86 | **Alkaloid** |
| Alkaloid_17 | 1796 | 0.43±0.42 | **Alkaloid** |
| Alkaloid_18 | 1806 | 0.23±0.05 | **Alkaloid** |
| Alkaloid_19 | 1810 | 0.36±0.31 | **Alkaloid** |
| Alkaloid_20 | 1817 | 0.40±0.09 | **Alkaloid** |
| Alkaloid_21 | 1828 | 0.47±0.53 | **Alkaloid** |
| Actinomycin C2 derivative_1 | 1830 | 0.74±0.83 | **Antibiotic** |
| Pentadecanoic acid_1 | 1845 | 0.18±0.08 | **Acid** |
| Alkaloid_22 | 1861 | 0.60±0.37 | **Alkaloid** |
| Alkaloid_23 | 1864 | 0.22±0.19 | **Alkaloid** |
| Alkaloid_24 | 1870 | 0.29±0.30 | **Alkaloid** |
| Pentadecanoic acid_2 | 1875 | 0.50±0.17 | **Acid** |
| Alkaloid_25 | 1894 | 0.62±0.30 | **Alkaloid** |
| Alkaloid_26 | 1898 | 0.13±0.13 | **Alkaloid** |
| Hydrocarbon | 1930 | 0.60±0.56 | CHC |
| Alkaloid_27 | 1932 | 0.43±0.22 | **Alkaloid** |
| Alkaloid_28 | 1946 | 1.79±0.75 | **Alkaloid** |
| Alkaloid_29 | 1949 | 0.62±0.54 | **Alkaloid** |
| Hexadecenoic acid_1 | 1953 | 0.62±0.69 | **Acid** |
| Hexadecanoic acid_2 | 1986 | 17.65±0.80 | **Acid** |
| Unidentified_21 | 1999 | 0.12±0.12 | NA |
| Heptadecenoic acid_1 | 2050 | 0.32±0.06 | Acid |
| Alkaloid_30 | 2054 | 0.59±0.16 | **Alkaloid** |
| Heptadecanoic acid_2 | 2068 | 0.27±0.04 | **Acid** |
| Octadecanol | 2085 | 0.23±0.17 | Alcohol |
| Octadecadienoic acid methyl ester | 2096 | 0.67±0.15 | Ester |
| Unidentified_22 | 2113 | 0.30±0.05 | NA |
| Octadecanoic acid methyl ester | 2129 | 0.41±0.05 | Ester |
| Octadecenoic acid_1 | 2152 | 21.44±9.09 | **Acid** |
| Octadecanoic acid_2 | 2176 | 7.80±1.04 | **Acid** |
| Actinomycin C2 derivative_2 | 2237 | 0.15±0.13 | **Antibiotic** |
| Fatty acid methyl ester | 2261 | 0.28±0.03 | Ester |
| Eicosanol | 2286 | 0.24±0.02 | Alcohol |
| 2-(8Z)-8-Heptadecen-1-yl-4,5-dihydro-oxazole | 2319 | 0.17±0.15 | |
| Fatty acid ester_1 | 2358 | 0.68±0.13 | Ester |
| Octadecenamide | 2364 | 0.95±0.20 | Amide |
| Octadecanamide | 2389 | 0.25±0.04 | Amide |
| Unidentified_23 | 2414 | 0.18±0.07 | NA |
| N,N-Dimethyl-octadecenamide | 2434 | 0.23±0.06 | Amide |
| 2-(Dimethylamino)ethyl-octadecadienoate | 2456 | 0.34±0.08 | |
| 2-(Dimethylamino)ethyl-octadecenoate | 2461 | 0.65±0.12 | |
| Hexadecanoic acid, 2-hydroxy-1-(hydroxymethyl)ethyl ester | 2509 | 0.20±0.09 | Ester |
| Fatty acid ester_2 | 2543 | 0.16±0.07 | Ester |
| Fatty acid ester_3 | 2565 | 0.22±0.18 | Ester |
| Dodecanoic acid, hexadecyl ester | 2954 | 0.21±0.06 | Ester |
| Fatty acid ester_4 | 3126 | 0.21±0.07 | Ester |

**Supplemental Table 13 | Chemical compounds in the metapleural gland.** Table includes information on compounds found in the MG samples (Extended Data Fig. 9). Chemical groups written in bold are known to have antimicrobial effects. Six MG were pooled per sample run in the GC-MS-TD (*n*=3).

**Supplemental Movie 1 | Example of wound care performed with metapleural gland secretions collected from the gland of the individual providing care.** The infected ant is marked in white. We first observe wound care by the nursing ant, followed by the collection of metapleural gland secretions using the forelegs to reach the gland and mouth and finally application of metapleural gland secretions on the wound.

**Supplemental Movie 2 | Example of wound care performed with metapleural gland secretions collected from the gland of the injured individual**. The infected ant is marked in white and red. We first observe wound care by the nursing ant, followed by the collection of metapleural gland secretions of the injured individual using its mouthparts to reach the gland and finally application of metapleural gland secretions on the wound.
